# Neural circuitry of a polycystin-mediated hydrodynamic startle response for predator avoidance

**DOI:** 10.1101/255570

**Authors:** Luis A. Bezares-Calderón, Jürgen Berger, Sanja Jasek, Csaba Verasztó, Sara Mendes, Martin Gühmann, Rodrigo Almeda, Réza Shahidi, Gáspár Jékely

## Abstract

Startle responses triggered by aversive stimuli including predators are widespread across animals. These coordinated whole-body actions require the rapid and simultaneous activation of a large number of muscles. Here we study a startle response in a planktonic larva to understand the whole-body circuit implementation of the behavior. Upon encountering water vibrations, larvae of the annelid *Platynereis* close their locomotor cilia and simultaneously contract the body and raise the parapodia. The startle response is mediated by collar receptor neurons expressing the polycystins PKD1-1 and PKD2-1. CRISPR-generated *PKD1-1* and *PKD2-1* mutant larvae do not startle and fall prey to a copepod predator at a higher rate. Reconstruction of the whole-body connectome of the collar-receptor-cell circuitry revealed converging feedforward circuits to the ciliary bands and muscles. The wiring diagram suggests circuit mechanisms for the intersegmental and left-right coordination of the response. Our results reveal how polycystin-mediated mechanosensation can trigger a coordinated whole-body effector response involved in predator avoidance.

**Short Summary:** The neuronal circuitry of the *Platynereis* startle response links polycystin-dependent hydrodynamic sensors to muscle and ciliary effector cells

## Introduction

Approaching predators or other threatening stimuli often elicit prey escape or startle responses characterized by rapid whole-body locomotory actions (Bullock 1984). Startle responses require the simultaneous activation of a large number of muscles in a coordinated manner with a short latency (Eaton et al. 1977). Previous work on the neural mechanisms of the startle response in various animals including crayfish, the jellyfish *Aglantha digitale*, lampreys, fish, and amphibian tadpoles have uncovered commonalities in the underlying neuronal circuitry. These include giant premotor command neurons such as the giant fibers in crayfish (Edwards et al. 1999), the giant motor axons in *A. digitale* (Roberts and Mackie 1980), the Mauthner-cells in fish and lampreys (Korn and Faber 2005), and a Mauthner-cell-like system in larval *Ciona intestinalis* (Ryan et al. 2017), a planktonic chordate. Another common motif in startle circuits is convergent excitation. In crayfish and fish, the command neurons receive convergent input from multiple sensory afferents and excitatory feedforward interneurons (Lacoste et al. 2015; Zucker 1972). In *Drosophila* larvae, a combination of mechanosensory and nociceptive cues synergistically activates a faster mode of escape locomotion (Ohyama et al. 2015). The motor programs during a startle response can also differ depending on the location of the stimulus due to differences in connectivity downstream of different sensory fields (Edwards et al. 1999; Takagi et al. 2017). Physiological and anatomical studies have found that startle circuits consist of diverse neuron types. In jellyfish, up to a dozen distinct neuron types with dedicated functions may be involved (Mackie and Meech 1995a; Mackie and Meech 1995b). In crayfish and fish, the command neuron circuitry contains elaborate networks of pre‐ and postsynaptic neurons (Edwards et al. 1999; Korn and Faber 2005). However, we lack a complete, synapse level description of the whole-body circuitry for a behaviorally characterized startle response.

## A hydrodynamic startle response in *Platynereis* larvae

Nectochaete larvae of the marine annelid *Platynereis dumerilii* (Figure 1A) are planktonic and swim with beating cilia arranged into segmental bands (trochs). The larvae have three main trunk segments, each with two pairs of parapodia endowed with spiny chaetae and a complex musculature (Fischer et al. 2010). Upon disturbances to the surrounding water, freely swimming larvae abruptly stop swimming, contract the body and simultaneously elevate all parapodia (Figure 1B-C). This response is followed by a slower recovery phase (Suppl. Figure 1, Video 1). Similar responses have been reported in other planktonic animals (Mackie et al. 1976), and have been suggested to defend against predators (Pennington and Chia 1984). Defense strategies in invertebrate planktonic larvae may play an important role to reduce mortality during their planktonic development and consequently, ensure successful larval dispersion and recruitment. Here we take advantage of the functional tools available for *Platynereis* (Verasztó et al. 2017; Randel et al. 2014; Randel et al. 2015) to dissect the molecular, neuronal and behavioral bases of the startle response.

**Figure 1:**
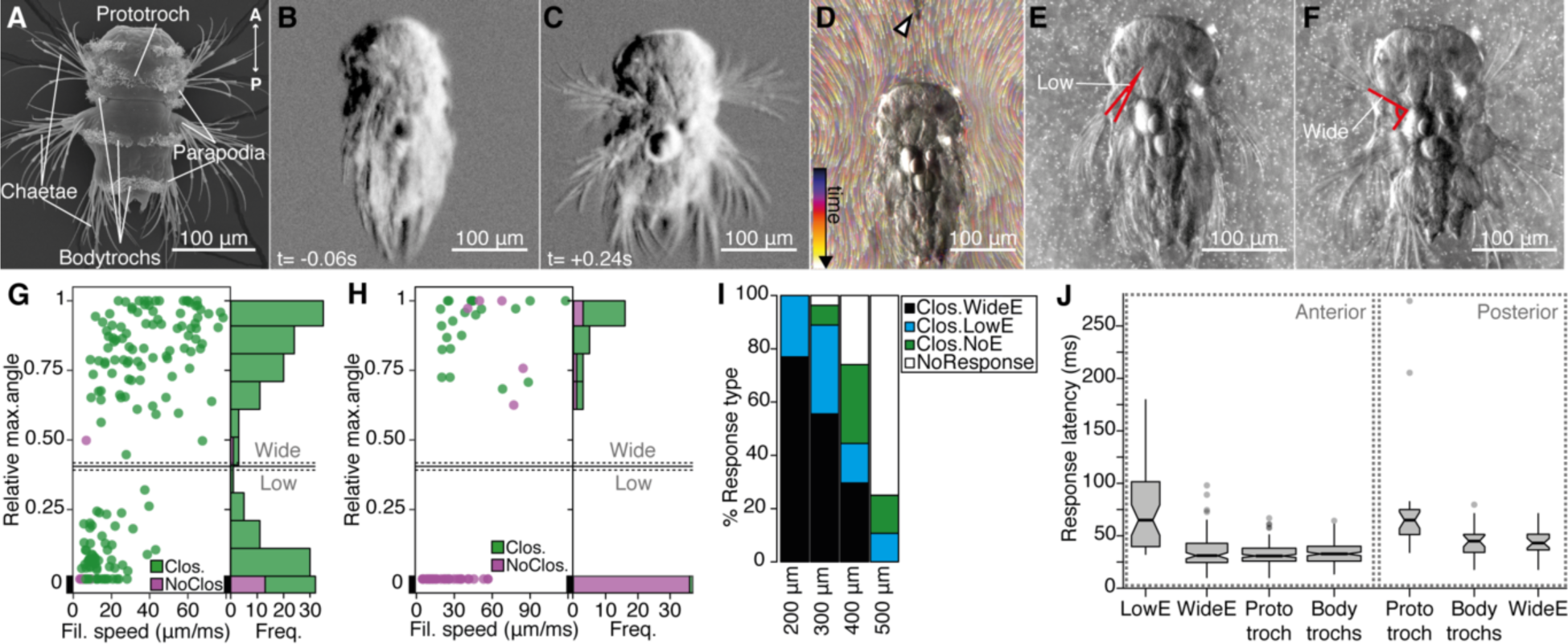
The startle response in Platynereis larvae is driven by hydrodynamic disturbances. (**A**) SEM micrograph of a 3-day-old nectochaete larva, dorsal view. (**B-C**) Snapshots of Video 1 60 ms before (**B**) and 240 ms after (**C**) a vibration stimulus. (**D**) Trunk-tethered larva (arrowhead points to the filament used for stimulation) engaging in (fictive) swimming. Fluorescent beads are color coded by frame. (**E-F**) Examples of low-angle (**E**) or wide-angle (**F**) parapodial elevation upon anterior stimulation (angles measured are outlined in red). (**G-H**) Relative parapodial elevation angle (0, no elevation; 1, maximum elevation) as a function of filament speed. The stimulation filament was placed 100 µm from the head (G) or pygidium (H). Dots are color coded by ciliary band (prototroch) closure. A stacked histogram is shown to the right of each scatter plot. A finite Mixture Model-based cutoff (0.4, solid horizontal lines) splits the two main populations of the histogram in (G, H), dashed horizontal lines: 95% CI. (**I**) Stacked barplot of startle response profiles upon stimulation with a filament placed at varying distances from the head. (**J**) Latency distributions for the onset of each response upon anterior (left) or posterior (right) stimulation. Notch in boxplots displays the 95% CI around the median. Boxwidth proportional to 
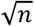
. n=9 larvae in panels **G** to **J**, each tested multiple times.

We first analyzed the kinematics of the startle response. We recorded trunk-tethered larvae at 230-350 frames per second during fictive swimming and stimulated them with a vibrating tungsten filament placed at a distance from the larvae to trigger startle responses (Figure 1D and Video 2). We visualized the water flow with fluorescent microbeads. When we stimulated larvae from the anterior, they closed their beating cilia and elevated the parapodia. Both aspects of the response were modulated as a function of filament speed (Figure 1G). The main ciliary band (prototroch) closed in all trials except at the lowest filament speeds tested (Figure 1G). The closures extended to all locomotor ciliary bands (here referred to as bodytrochs) (Video 3). Although parapodial elevation was triggered with some of the lowest filament speeds tested, it was only consistently observed above 28 µm/ms. The extent of the elevation of parapodia was dependent on the speed of the filament and displayed a bimodal distribution (Figure 1G). We split the distribution into low-angle and wide-angle elevation responses (LowE and WideE, respectively) by a cutoff obtained from adjusting the data to a finite mixture model (Figure 1E-G, Video 3). LowE responses were more common at low-to-moderate speeds, while WideE responses were seen across a broad stimulus-intensity spectrum (Figure 1G, Suppl. Figure 1). WideE responses had a shorter latency and the maximum parapodial elevation was achieved sooner than LowE responses (Figure 1J, and data not shown). Ciliary arrests and WideE responses occurred in close succession to each other (Suppl. Figure 1). As we moved the filament further away from the head, the response profile shifted to greater filament speeds (Figure 1I), showing that the response is triggered by near-field hydrodynamic disturbances.

Next we stimulated larvae from the posterior end (pygidium) from a 100 µm distance. Posterior stimulation triggered startle behaviors with a markedly different response profile compared to anterior stimuli (Figure 1H). First, animals were less sensitive to posterior stimulation as only filament speeds >19 µm/ms triggered a response. Second, we only observed WideE and no LowE responses. The WideE responses were often not accompanied by ciliary closures, but were as fast as WideE responses triggered from anterior stimulation (Suppl. Figure 1). Strikingly, we only very rarely observed ciliary closures not accompanied by a WideE response. However, ciliary closures and the WideE response were temporarily uncoupled (ciliary closures median latency: 65 ms (prototroch) or 45.6 ms (bodytrochs); WideE response median latency: 42 ms)(Figure 1J, K).

During WideE responses the three chaeta-bearing segments and the left and right body sides were highly coordinated. As already apparent from freely swimming larvae (Figure 1C) the extent of the elevation was bilaterally symmetric in all responses observed (Figure 1F shows a representative example). The delay values between the onset of elevation of the first pair of left and right parapodia were mostly below our recording speed limit (Suppl. Figure 1). Parapodia in different segments, but on the same body side rose simultaneously or one frame apart from each other (Suppl. Figure 1).

We thus characterized a rapid, left-right symmetric, segmentally coordinated, stereotypical whole-body startle response triggered by hydrodynamic stimuli with differences in response profile to anterior and posterior stimuli.

## Collar-receptor neurons respond to water-borne vibrations

To understand the neuronal mechanism of the startle response, we searched for candidate mechanosensory neurons in *Platynereis* larvae. The most likely candidates to sense flow-driven strain and bending are neurons with ciliary structures penetrating the cuticular surface (Mackie et al. 1976). Using a combination of scanning (SEM) and serial-section transmission electron microscopy (ssTEM), we identified and mapped all penetrating uniciliated, biciliated and multiciliated sensory neurons across the whole body of nectochaete larvae (Figure 2 and Suppl. Figure 2). Since nectochaete larvae have a stereotypical morphology, the same cells could be reliably identified in different larvae by SEM and in the ssTEM dataset.

**Figure 2:**
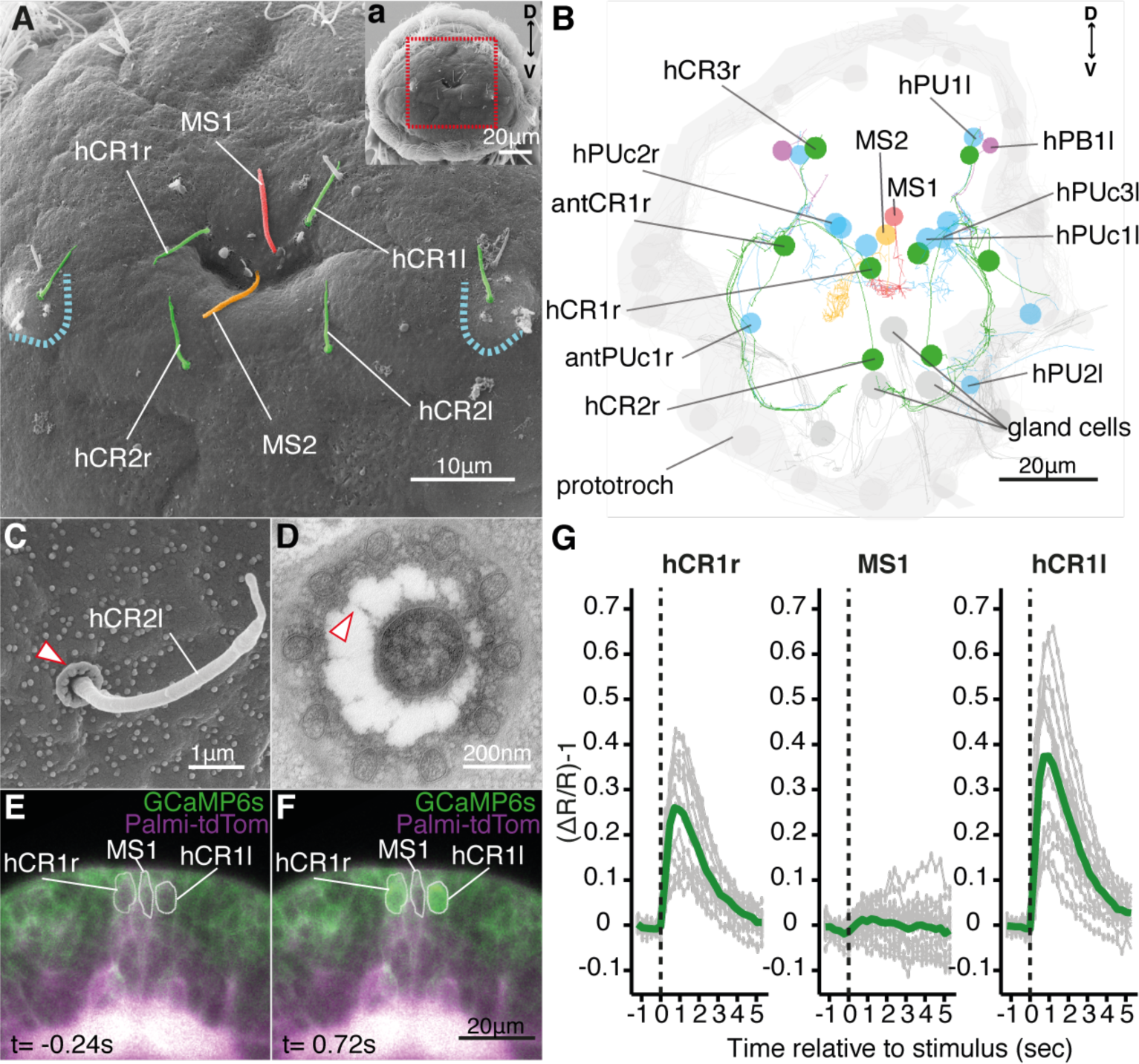
Head CR neurons are hydrodynamic receptors. (**A**) Close-up view of the head of a nectochaete larva (region outlined in **a**). Developing antennae with a CR cilium (green) are outlined in blue. (**B**) Reconstruction of penetrating ciliated sensory neurons in the head from a serial transmission electron microscopy volume. Color code as in (A). Additional penetrating uniciliated (PUc and PU) or biciliated (PB) neurons are shown in blue and magenta, respectively. One of each bilateral pair is labeled. (**C**) SEM of a CR sensory cilium (hCR2l), collar of microvilli protrudes out of the cuticle (arrowhead). (**D**) TEM cross-section of a CR sensory cilium with a collar of 10 thick microvilli and fibers connecting the cilium and the microvilli (arrowhead). (**E, F**) Snapshots from Video 5 showing a larva injected with *GCaMP6s-P2A-Palmi-tdTomato* mRNA 0.24 sec prior (**E**) or 0.72 sec after (**F**) stimulation with a vibrating filament (ventral view). The ROIs used for fluorescence quantification are outlined in white. (**G**) Mean ratiometric fluorescence changes (green traces) of hCR1l, hCR1r and MS1 neurons upon stimulation with a vibrating filament placed 50 µm anterior to the head. 15 (hCR1r), 18 (MS1) and 16 (hCR1l) measurements (gray traces) from 5 animals are shown. The traces were aligned relative to the stimulus start (t=0).

We narrowed down the list of candidate hydrodynamic receptors to those penetrating ciliated cells found on the head of the animal, as this region was the most sensitive to mechanosensory stimuli (Figure 1). We found different candidate cell types (Figure 2A-B), including a group of cells very similar to the collar receptor cells (CRs) previously identified in other polychaetes, (Budelmann 1989; Schlawny et al. 1991; Windoffer and Westheide 1988; Purschke 2005; Purschke et al. 2016), earthworms (Knapp and Mill 1971) and leeches (called ‘S hairs’) (Phillips and Friesen 1982). In leeches, these cells were suggested to be involved in the detection of water movement (Friesen 1981). The CR neurons in *Platynereis* have a single non-motile cilium with a 9×2+2 microtubule doublet pattern and a symmetric collar of 10 thick microvilli surrounding the cilium (Figure 2C-D). Each microvillus has a dense region in the inner side and is connected by thin fibers to the cilium (Figure 2D). The collar and the cilium penetrate the cuticle, thus these cells can also be identified in SEM samples. CR neurons also occur in other regions of the nectochaete larva, either individually or in clusters as parts of developing organs (Figure 2A, B, and Suppl. Figure 2).

To determine if the head CR neurons are hydrodynamic mechanoreceptors, we recorded the activity of a set of clearly identifiable head CR neurons (hCR1 and hCR2) by calcium imaging (Figure 2E, Suppl. Figure 3 and Video 4). We also imaged another distinct type of collared uniciliated penetrating neuron called MS1 located between the two hCR1 cells (Figure 2A-B and Suppl. Figure 2). The cell bodies of hCR1 and MS1 neurons lie in the same focal plane and could be recorded simultaneously (Figure 2E, F, Suppl. Figure 3). When we stimulated larvae with a vibrating filament, the cilia of the hCR1 cells deflected and GCaMP6s fluorescence increased in the cell bodies and in the cilia (Figure 2E-G, Video 5 and Video 6). Although the cilium of MS1 was also deflected, the fluorescence did not increase in this cell (Figure 2G, and Video 6). hCR2 neurons were also activated by the stimulus (Suppl. Figure 3). These results show that head CR neurons detect hydrodynamic vibrations and could thus trigger the startle response.

## CR neurons express TRPP and polycystin channels

To investigate the mechanism of hydrodynamic reception, we searched for mechanosensory markers specifically expressed in CR neurons. We found that a *Platynereis* ortholog of the TRPP/Polycystin-2 channel, *PKD2-1*, (Suppl. Figure 4) was expressed in CR neurons, as demonstrated by *in situ* hybridisation and a transgenic reporter construct (Figure 3A, B). *PKD2-1* was also expressed in two other types of penetrating uni‐ and biciliated sensory neurons (PU and PB neurons, respectively) in the head, trunk, and pygidium, including an unpaired biciliated sensory cell in the pygidium (*pygPB^unp^*) (formerly *pygPB^bicil^* (Shahidi et al. 2015; Verasztó et al. 2017)) (Suppl. Figure 5). Another gene in *Platynereis*, *PKD1-1*, was expressed only in CR neurons and in the *pygPB^unp^* neuron at the nectochaete stage (Figure 3C, D and Suppl. Figure 6). *PKD1-1* belongs to the TRPP-related family PKD1, and it defines a novel invertebrate family that is paralogous to the more widespread Polycystin-1/PKD1L1 families (Suppl. Figure 4).

**Figure 3:**
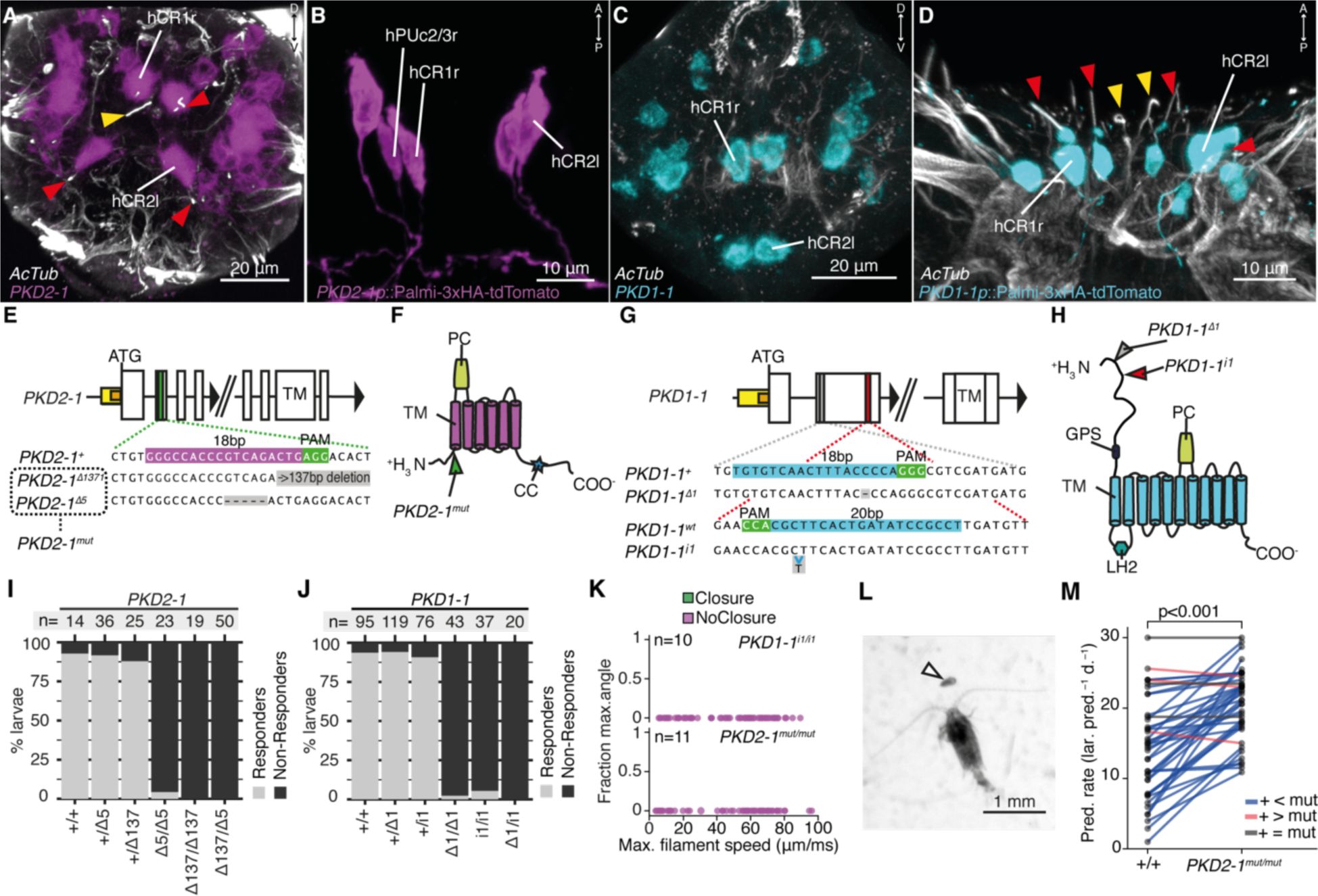
CR neurons express PKD1-1 and PKD2-1, and are required for the startle response. (**A** and **C**) PKD2-1 (A) or PKD1-1 (C) head gene expression revealed by in situ hybridization; apical views. (**B** and **D**) Immunostaining of HA-Palmi-tdTomato expressed under the PKD2-1 (B) or the PKD1-1 (D) promoter; ventral views. (**E** and **G**) PKD2-1 (E) and PKD1-1 (G) genomic loci and close-up regions showing wildtype (+) sequences targeted with CRISPR/Cas9. The mutant alleles generated are also shown. White, yellow and orange boxes represent exons, promoters, and 5’UTRs, respectively. (**F** and **H**) PKD2-1 (F) or PKD1-1 (H) protein architecture. Main conserved domains are labeled. Mutation sites are indicated by arrowheads. (**I** and **J**) Stacked barplots of PKD2-1 (I) or PKD1-1 (J) mutant and wildtype (+/+) homozygote, heterozygote or trans-heterozygote larvae (%) startle response to touch stimulus. (**K**) Parapodial elevation angles of PKD1-1^i1/i1^ or PKD2-1^mut/mut^ larvae as a function of filament speed. Stimulation filament ca. 100 µm from the head. (**L**) Adult Centropages typicus female approaching a nectochaete Platynereis larva (arrowhead). (**M**) Predation on wildtype (+/+) and PKD2-1^mut/mut^ larvae by C. typicus. Paired values are joined by blue, gray or red lines if predation rates were higher, equal or lower in mutant than in wildtype larvae, respectively; data from 42 trials with 12 batches. One-sided exact Wilcoxon-Pratt signed rank test, P < 0.001. Arrowheads in A-D point to CR cilia (red) or to MS1/2 cilia (yellow).

## *PKD1-1* and *PKD2-1* are required for the startle response

The co-expression of *PKD1-1* and *PKD2-1* in CR neurons and the known function of their homologs in mechanosensation (Sharif-Naeini et al. 2009) suggested that polycystins participate in the mechanotransduction cascade in CR neurons. To test this, we generated mutant lines with the CRISPR/Cas-9 system (Lin et al. 2014). We recovered multiple deletion alleles in both *PKD1-1* and *PKD2-1* with frameshift mutations and premature STOP codons leading to protein products predicted to lack most functional domains (Figure 3E-H). Homozygote and trans-heterozygote larvae for either *PKD1-1* or *PKD2-1* alleles were viable and fertile and had normal CR neuron morphology (Suppl. Figure 7). These larvae, however, failed to startle upon touching their head, a stimulus that triggered the startle response in most wild type and heterozygote larvae (Figure 3I-J). We also tested the responses of tethered *PKD1-1* and *PKD2-1* mutant larvae stimulated from the anterior or the posterior end with a vibrating filament (Figure 3K, Suppl. Figure 7 and Video 7). All aspects of the startle response, including ciliary arrests and parapodial extensions were absent from mutant larvae for either of the genes across the whole range of filament speeds tested. This demonstrates that the neuronal expression of *PKD1-1* and *PKD2-1* channels is essential for the hydrodynamic startle response in *Platynereis* larvae.

The absence of a startle response in the otherwise normal polycystin mutants allowed us to test whether the startle response plays a role in prey-predator interactions, as suggested for other planktonic larvae that have a similar response (Pennington and Chia 1984; Thiel et al. 2017). We exposed *Platynereis* larvae to *Centropages typicus*, a predatory copepod which detects its prey by the hydromechanical signals it generates (Calbet et al. 2007; Kerfoot 1978)(Figure 3L). We incubated in the same container an equal amount of *PKD2-1^mut/mut^* and age-matched wildtype larvae with *C. typicus* and quantified the number of survivors of each genotype after 12h or 24h. We also incubated larvae without copepods to rule out differences in mortality rates not related to predation (Table 1). In most experiments, *PKD2-1^mut/mut^* larvae were predated more than their wildtype counterparts (*P*< 0.001, one-sided exact Wilcoxon-Pratt signed rank test, Figure 3M). As wild type larvae showed the startle response upon copepod attacks (Video 8), the difference in predation rates is likely due to the absence of a startle response in *PKD2-1^mut/mut^* larvae.

## Whole-body wiring diagram of the startle response

The hydrodynamic sensitivity of the head CR neurons (Figure 2), their expression of *PKD* genes, and the requirement of these genes for the startle response (Figure 3) argue that the CRs are the main sensory receptors that trigger the startle response. To investigate the neuronal mechanisms behind the response, we mapped the wiring diagram downstream of all CRs in a whole-body ssTEM dataset. We identified all the direct postsynaptic partners (> *2* synapses from > 1 CRs) of CRs in the larva (Table 2 and 3). We only analyzed the most direct neural paths from CRs to the ciliated and muscle effectors, as the short latency of the response suggests that there are few intervening synapses. The head and pygidial CRs project along the ventral nerve cord (VNC) posteriorly and anteriorly, respectively (Figure 4A and Suppl. Figure 2), and form *en passant* synapses with distinct types of interneurons (INs) and motorneurons (MNs) (Figure 4B-F, Suppl. Figure 8 and Video 9). The IN layer targets several muscle-motor and ciliomotor neurons forming a wiring diagram that can explain the main aspects of the startle response.

**Figure 4:**
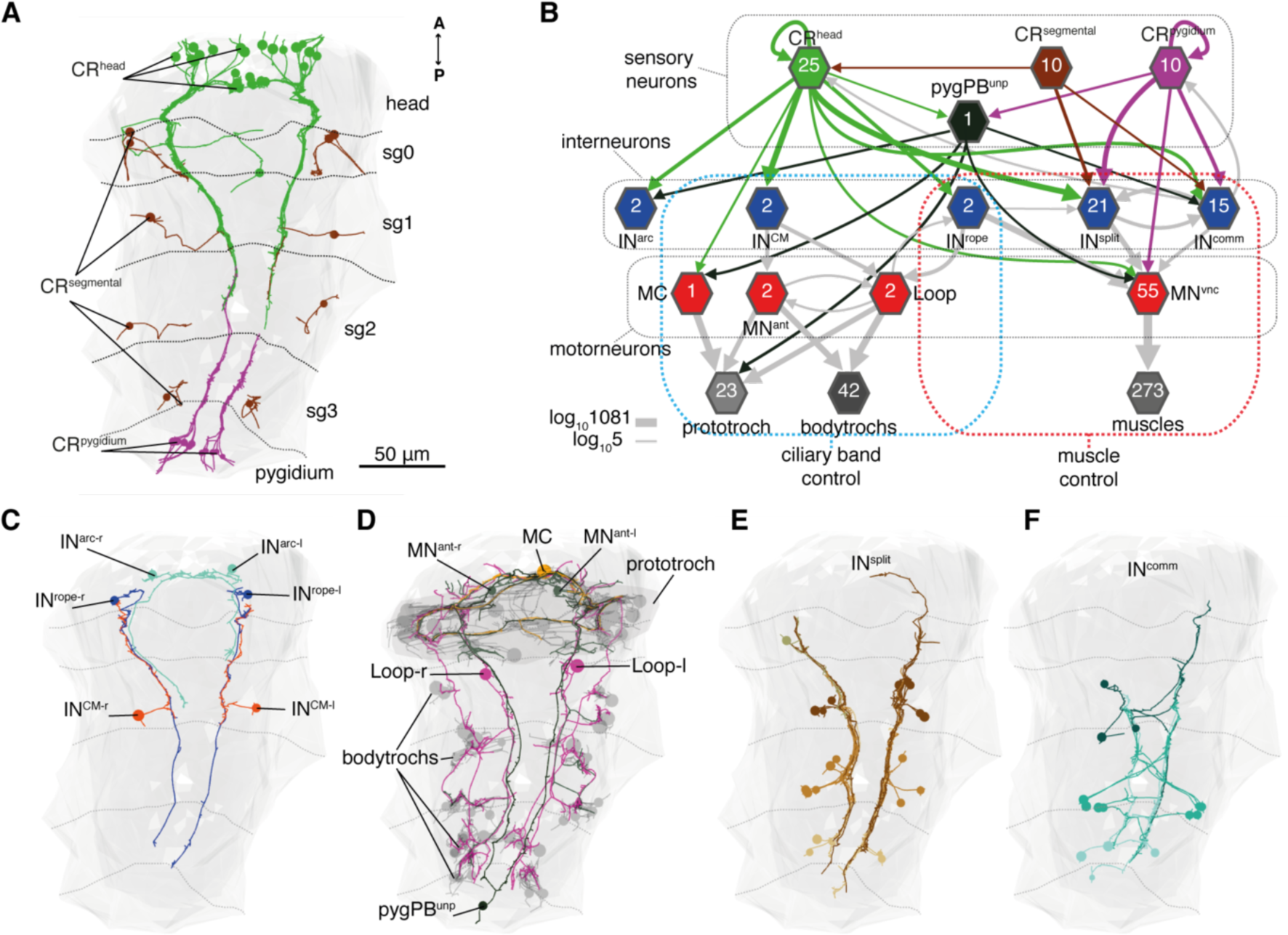
Wiring diagram of CR neurons. (**A**) All CRs in the nectochaete larva reconstructed from an EM volume. Head, segmental and pygidial groups are differently colored. Segment (sg) boundaries are indicated by dotted lines. (**B**) Sensory-motor network from CR neurons to muscles and locomotory cilia. Hexagons and arrows represent neuron groups and their synaptic connections, respectively. Numbers inside hexagons indicate the number of neurons grouped in each node. Interactions with less than 5 synapses were filtered out in this display. (**C**) EM reconstruction of Rope, Arc and CM neurons. (**D**) Ciliomotor neurons targeted by CRs or by IN^CM^ (**E**) Ipsilateral split neurons (IN^split^) (**F**) Commisural interneurons (IN^comm^). Ventral view in A, C-F. Segment boundaries as defined in A are overlaid in C-F.

The head CRs directly synapse on the MC cell, a cholinergic motorneuron that induces ciliary closures of the prototroch ciliary band (Verasztó et al. 2017) (Figure 4B). This can partly explain that ciliary closures occur soon after anterior stimulation. The head but not the pygidial CRs also synapse extensively on CM interneurons (IN^CM^), a bilateral pair of ipsilaterally projecting pseudo-unipolar neurons with a soma in the 1^st^ segment (Figure 4B-C). The CMs are presynaptic to the Loop and MN^ant^ ciliomotor neurons controlling the prototroch and all bodytrochs (Verasztó et al. 2017)(Figure 4B). The activation of the ciliomotor through the CMs would close cilia on all the bodytrochs and the prototroch in a coordinated fashion and independent of muscle contraction. The head CRs also target another pair of interneurons, the Rope interneurons (IN^rope^). These neurons have their soma in the head and have ipsilateral descending projections that span the entire VNC (Figure 4C). The Rope neurons make synaptic contacts with the Loop neurons (Figure 4B), thus providing a parallel pathway to ciliary control. These feed-forward circuits from CRs to ciliated cells likely drive the closure of all locomotor cilia in a coordinated fashion upon CR activation (Figure 1G). Finally, both head and pygidial CRs target the sensorymotor neuron pygPB^unp^, which may modulate the prototroch ciliary band system (Verasztó et al. 2017).

CRs also feed into distinct muscle-motor pathways (Figure 4B). Both head and pygidial CRs directly synapse on the Crab motorneurons (MN^crab^), a unique set of decusating VNC motorneurons that innervate a variety of trunk muscles (Figure 4B, Suppl. Figure 8-9). A second pathway is through the Rope interneurons (Figure 4B). The descending projections of Rope neurons target several segmentally arranged ipsi‐ and contralaterally projecting VNC motorneurons that innervate distinct sets of somatic muscles (longitudinal, oblique, transverse, and parapodial) in all three segments (Figure 5A-B, Suppl. Figure 8-9). The Rope neurons in *Platynereis* are reminiscent of descending premotor interneurons in other startle circuits, including the M-cells in vertebrates, the GF cells in *Drosophila*, or the LG cells in crayfish (Hale et al. 2016). The Rope interneurons could thus have the analogous function of activating trunk motorneurons across segments that in turn would coordinately contract the muscles during the startle response. The involvement of Rope interneurons in both ciliary and muscular control suggests that they coordinate ciliary closures and whole-body muscle contractions upon anterior stimulation (Figure 1J and Suppl. Figure 1).

**Figure 5:**
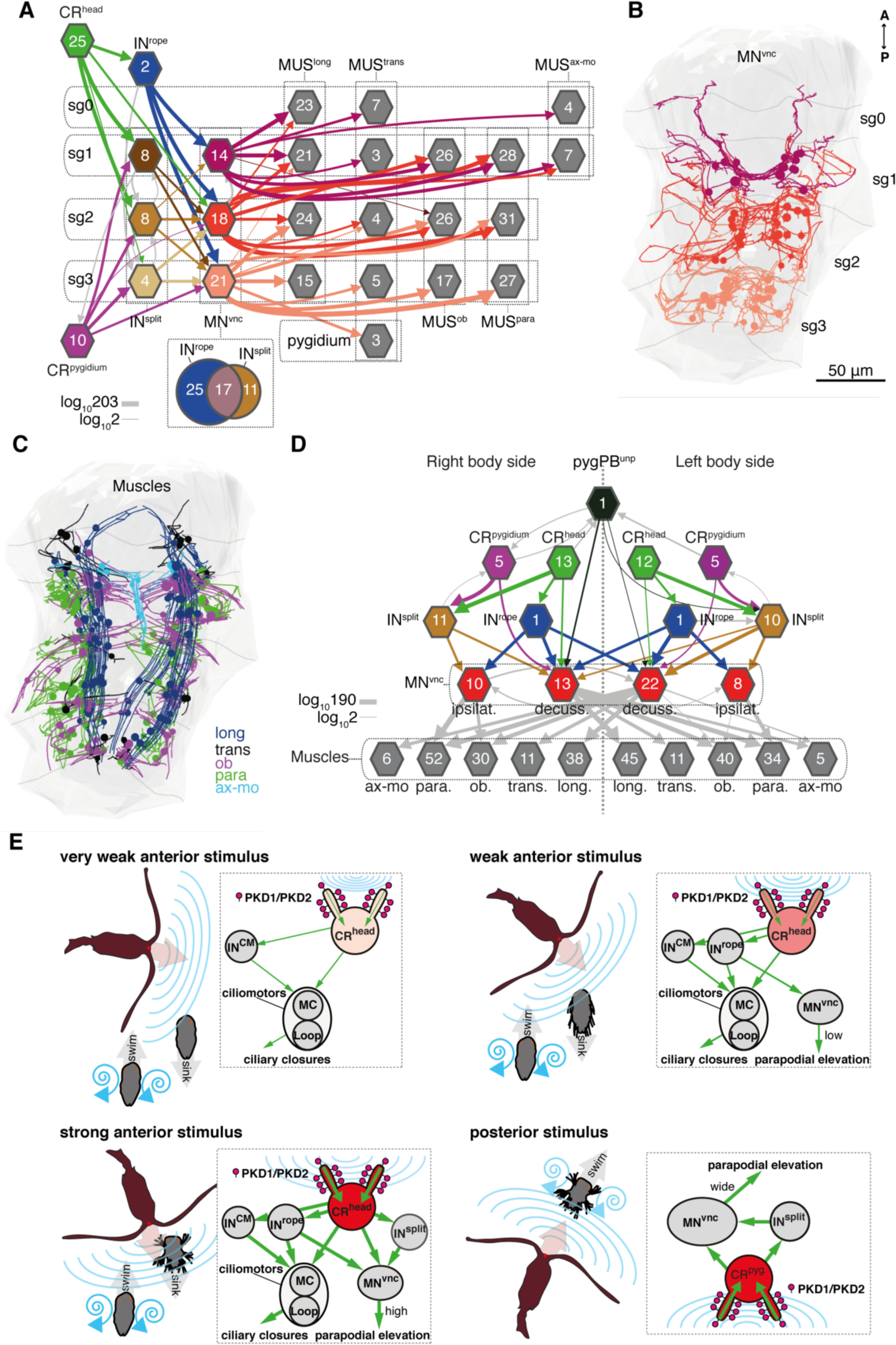
Bilateral and segmental coordination motifs in the startle circuit. (**A**) Muscle network downstream of Rope and Split interneurons sorted by segmental location and muscle type. Arrows represent synaptic connections and are colored based on the source neuronal group (represented by hexagons). (**B**) VNC motorneurons (MN^vnc^) targeted by Rope and Split interneurons colored by segmental location. (**C**) Muscles targeted by MN^vnc^ neurons colored by type (long, longitudinal; trans, transvers; ob, oblique; para, parapodial; axmo, axochord-mouth). (**D**) Rope and Split muscle network sorted by body side. (**E**) Summary schematics of the neural implementations behind the larval response in different predator-prey scenarios (right and left, respectively). Numbers inside hexagons in A and D indicate the number of neurons grouped in each node. Segment (sg) boundaries are indicated by dotted lines in A and C. Interactions with less than 2 synapses were filtered out in B and D.

An alternative muscle-motor pathway is through the Split interneurons (IN^split^), a group of segmentally arranged cells with a pseudounipolar morphology (similar to CMs)(Figure 4D and Suppl. Figure 8). The Split neurons connect to a set of VNC motorneurons in all segments that partially overlaps with the targets of the Rope interneurons (Figure 5A). The Split interneurons are the main targets of non-head CRs, suggesting that this pathway triggers muscle contractions independently of ciliary closures upon pygidium CR activation (Figure 4B).

CRs also synapse on a heterogenous group of ascending and descending trunk commissural interneurons (IN^comm^) (Figure 4E and Suppl. Figure 8). In contrast to the other interneuron types, we could not find bilateral pairs of IN^comm^ targeted by left and right CRs. Some of the IN^comm^ cells synapse on contralateral interneurons and motorneurons in the startle network and also feed back to the CRs (Figure 4B). These interneurons may thus coordinate both body sides during the response.

The VNC motorneurons targeted by the Rope and Split cells belong to at least ten morphologically distinct groups with segmentally iterated and left-right symmetric members (Figure 5A, B, D, Suppl. Figure 9). The motorneurons have both ipsilateral and decussating types. Together, these motorneurons innervate distinct muscle types that are recruited during a startle response, as assessed by calcium imaging (Figure 5C and Video 10).

A hallmark of the startle response is its left-right symmetry, both in timing and in the extent of muscle contraction (Figure 1 and Suppl. Figure 1). The proximity of the sensory cilia of all head and pygidium CRs suggests that both the left and right sides are stimulated in most cases, as observed during calcium imaging of hCR1 cells (Figure 2E-G). This could partly explain the bilateral symmetry of the response. However, there are also circuit motifs that likely ensure the left-right coordination of muscle and ciliary effectors. The pair of Rope neurons connect to decussating motorneurons on the same and opposite sides and both CMs connect to the ciliomotor MN^ant^ cells of both body sides (Figure 5D).

## Discussion

CR neurons constitute a newly identified mechanoreceptor cell type with a conserved morphology in annelids and with similarities to hair cells in vertebrates. We uncovered an essential role of PKD1-1 and PKD2-1 proteins in the transduction of hydrodynamic signals in CRs in *Platynereis dumerilii*. The neural role of polycystins is poorly understood, although they have been linked to flow detection in non-neuronal contexts (Yoshiba et al. 2012; Nauli et al. 2003). PKD1 plays only a structural role in mammalian hair cells and does not have a mechanosensory function (Steigelman et al. 2011). In invertebrates, only *C. elegans* PKD1 and PKD2 are known to be required for a neuronal mechanosensory function (Barr and Sternberg 1999).

The coexpression of *PKD1-1* and *PKD2-1* in CRs in *Platynereis* suggests that their protein products form a complex as it is known for other polycystins (Hanaoka et al. 2000; Nauli et al. 2003). Given that mutating these genes eliminated the startle response, we could identify the CR neurons as the sensory cells that mediate the behavior.

Whole-body connectomics of CRs identified converging feed-forward circuits from the hydrodynamic sensory CRs to the ciliary bands and to trunk muscles. The wiring diagram can explain the graded tuning, directional sensitivity and whole-body coordination of the startle response.

There are several circuit motifs for intersegmental and left-right coordination. Rope interneurons synapse on segmentally iterated left and right motorneurons. The axons of pygidium CRs synapse on the Crab motorneurons. Spider motorneurons span two segments, the intersegmental Loop and MN^ant^ ciliomotors span the head and trunk (Verasztó et al. 2017).

The diverse interneuron pathways likely have different activation thresholds ensuring graded responses to varying stimulus strengths. Ciliary closures could be driven by low-threshold pre-ciliomotor CMs. Rope interneurons that connect to both muscle‐ and ciliomotors could trigger synchronised ciliary arrests and parapodial extensions upon intermediate or high intensity stimulation. The different response profiles to anterior and posterior stimuli agree with the differential connectivity of head and tail CRs. LowE responses only occurred upon head stimulation and must be mediated by head-CR-specific connectivity (most likely through the Ropes).

The *Platynereis* startle circuit resembles other startle circuits. Ropes are intersegmental premotor neurons similar to the Mauthner-cells and may be the precursors to the giant interneurons reported in adult *Platynereis* (Smith 1957). These giant cells target the 4B motorneurons, which have a similar morphology to the MN^crab^ neurons described here (Sigger and Dorsett 1986). The convergence of feed-forward motifs is also present in the *Platynereis* circuit, as in other startle circuits. All interneuron classes integrate CR input, and interneurons and CRs also converge on motorneurons. For example, the MN^crab^ neurons receive input from both Rope and Split interneurons as well as head and tail CRs.

We have analyzed an ecologically relevant whole-body startle behavior and the molecular machinery and neuronal circuitry underlying it (Figure 5E). The *Platynereis* startle response and the circuit mediating it are likely fine tuned to maximize the avoidance of predators by the planktonic larvae. Beating cilia generate a hydrodynamic signal that can be detected by predators (Kiørboe and Visser 1999). The closure of all cilia triggered by weak mechanical signals could help the larvae to avoid being detected. Mechanically induced ciliary arrests have been reported in other ciliated larvae (Mackie et al. 1976) and could be a widespread avoidance mechanism. Higher intensity stimulation by nearby predators could trigger chaetal extension for deterrence. Posterior stimulation often only induced chaetal extension while cilia continued beating, potentially increasing the chances of a forward escape (Figure 5E). The study of the *Platynereis* larva allowed us to analyze in a whole-body context the molecular, cellular and circuit mechanisms of an ecologically relevant behavior.

## Materials and Methods

### Animal culture and behavioral experiments

*Platynereis dumerilii* wild type (Tübingen) strain and the mutant lines derived from it were cultured in the laboratory as previously described (Fischer and Dorresteijn 2004). The nectochaete larval stage (approximately 72 hours post fertilization) was utilized for all experiments and raised at 18°C on a 16:8-h light/dark cycle in glass beakers. Filtered natural sea water (fNSW) was used for all the experiments.

Adult stages of the marine planktonic copepod *Centropages typicus* were used as rheotactic predators for wild type and *PKD2-1* mutant larvae. Copepods were supplied from a continuous culture at the National Institute for Aquatic Resources (Technical University of Denmark, DTU). Specimens of *C. typicus* were originally isolated from zooplankton samples collected in Gullmar fjord (Sweden) by vertical tows with plankton nets (500 µm mesh). Cultures of *C. typicus* were kept in 30 L plastic tanks with sterile-filtered seawater (FSW, salinity 32 ppt), gently aerated, at 16 ± 1 °C in dark. Copepods were fed *ad libitum* with a mix of phytoplankton (the cryptophyte *Rhodomonas sp.*, the diatom *Thalassiosira weissflogii* and the autotrophic dinoflagellates *Heterocapsa triquetra*, *Prorocentrum minimum* and *Gymnodinium sanguineum*) and with the heterotrophic dinoflagellate *Oxyrrhis marina*. Phytoplankton cultures were kept in exponential growth in B1 culture medium and maintained at 18°C and on a 12:12-h light/dark cycle in glass flasks. *O. marina* was fed the cryptophyte *Rhodomonas salina* and maintained at 18°C in 2-L glass bottles. Behavioral experiments were performed at room temperature unless otherwise indicated.

### Kinematics of startle behavior

Larvae were relaxed in 50-100 mM MgCl_2_ 10 min before tethering them to a Ø3.5 cm glass-bottom dish (HBST-3522, Willco Wells) with a non-toxic glue developed for *C. elegans* (Wormglu, GluStitch Inc). Care was taken to minimize or to avoid gluing ciliary bands, sensory cilia, parapodia, head and pygidium. Prior to the experiment, the glued larvae were assessed for the startle response with a gentle vibration to verify that relevant structures were unhindered and the animal was healthy and in a predominantly swimming mode. 1 µm multi-fluorescent beads (24062-5, Polysciences) were diluted in 5% BSA to 1:100, sonicated for 1 min and added to the glued larva preparation at a 1:10 dilution. The experiments were done in a final volume of 2.5 ml. Recordings were done with an AxioZoom V.16 (Carl Zeiss GmbH, Jena) and responses were recorded with a digital CMOS camera ORCA®-Flash-4.0 V2 (Hamamatsu). A HXP 200 fluorescence lamp at maximum level (using the Zeiss 45 mCherry filter) was switched on only during each recording (lasting max. 7 sec each) to visualize the beads.

To generate water-borne vibrations, a thin 3-5 cm tungsten needle (RS-6063, Roboz) was glued to a shaftless vibration motor (EXP-R25-390, Pololu), which was switched on for a defined time interval (1-35 ms) and induced the needle to vibrate. The motor was switched on and off with a custom script via an Arduino microcontroller (Arduino UNO R3,Arduino). The probe was positioned in focus at a defined distance from the larva with the manual micromanipulator US-3F (Narishige). A defined set of stimulation values were used for each of the animals tested, the order of the values was randomized for each larva. Between each stimulation attempt, the larva was left to rest for 1 min. Recordings were discarded if the larva was not in swimming mode while being stimulated. Stimulus start was defined from the videos by the onset of probe movement. The larvae were still alive and visibly healthy one day after being glued. All wild type larvae tested came from different batches. In the case of mutants, 10 larvae from two *PKD1-1^i1/i1^* batches and 11 larvae from three *PKD2-1^mut/mut^* batches were tested. Each tested larva was genotyped after the experiment.

The parapodial elevation angle was measured as the movement of the distal end of one of the parapodia in the first segment relative to its base that occurred from the rest position prior to stimulation to the maximal elevation achieved upon stimulation. The value was normalized to the maximum angle recorded for a given larva. Filament speed was calculated as the maximum displacement observed between consecutive frames normalized to the recording speed. Onset of ciliary band closures was defined as the moment after stimulation the flow of fluorescent beads stopped in the vicinity of the cilia. For bodytroch closures this was assessed from the posterior-most band (telotroch). Onset of parapodial elevation was defined as the moment after stimulation the first segment parapodium (from which the elevation angle was measured) started its angular elevation. The slow elevation speed for LowE responses in many cases made difficult to assess with precision the start of the elevation movement.

### Startle assay in freely swimming animals

4ml of phototactic nectochaete larve were transferred to a Ø5 cm glass-bottom dish (GWSB-5040, Willco Wells) with a vibrating shaftless motor glued to its bottom. The motor was activated for 100ms via an Arduino microcontroller with a custom-written script. The same microscopy equipment used for the kinematics experiments was used, but the recording speed was set to 15 fps. A long-pass filter was placed between the light source and the dish to minimize phototaxis. The speed of startled larvae was calculated as the Pitagorean distance over time unit, and the area was measured from the thresholded shape of the larva.

### Whole Mount *in situ* Hybridization and Immunochemistry

*In situ* hybridization probes for detecting *PKD1-1* and *PKD2-1* were synthesized from 1 Kb gene fragments subcloned into pCR™II vectors (K206001, ThermoFisher) (Table 4), or from an in-house EST vector library. Whole mount *in situ* hybridization was done as previously described (Conzelmann et al. 2011). *PKD1-1* and *PKD2-1* promoters (fragment sizes: 2.5 Kb and 1.5 Kb, respectively) were amplified and cloned upstream of *3xHA-Palmi-tdTomato* (Table 4). Larvae injected with promoter constructs (ca. 250 ng/µl) were analyzed for reporter expression at 3 days post fertilization using an AxioImager Z.1 fluorescence wide-field microscope (Carl Zeiss GmbH, Jena) and immediately fixed for immunostainings. The protocol followed for immunostaining of HA-tagged reporters was recently described (Verasztó et al. 2017). Imaging of specimens was performed in a LSM 780 NLO Confocal (Zeiss, Jena).

### Generation of *PKD1-1* and *PKD2-1* mutants with CRISPR/Cas9

The SpCas9-GFP ORF was kindly provided by Jennifer Doudna (Addgene plasmid #42234) and was subcloned into a custom-made plasmid construct tailored for enhanced mRNA expression in *Platynereis*. sgRNAs were designed using ZiFiT (Sander et al. 2007), and cloned into either the DR274 plasmid (kindly provided by Keith Joung, Addgene plasmid #42250) or pX335-U6-Chimeric-BB-CBh-hSpCas9n (D10A) (kindly provided by Feng Zhang, Addgene plasmid #42335). sgRNAs were synthesized from PCR templates (Table 4) with the MEGAshortscript T7 kit (AM1354, Ambion) and purified using the MEGAclear Transcription Clean-up kit (AM1908, Ambion). Zygotes were injected with a mix of 300 ng/µl SpCas9-GFP mRNA and 20 or 50 ng/µl sgRNA dissolved in DNAse/RNase-free water (10977-049,ThermoFisher). 1-day-old larvae were screened for green fluorescence at 1 day post fertilization and gDNA was extracted from single larvae in 4 µl QuickExtract (QE) Solution (QE09050,Epicentre). Mutations were detected by PCR and Sanger sequencing (Table 4). Adult (atokous) worms injected with effective sgRNAs were genotyped by clipping the pygidium following a previously published protocol (Bannister et al. 2014), but using 20 µl of QE solution for gDNA extraction. Worms carrying desired mutations were outcrossed to the wild type strain for the first two generations before using them for experiments.

### Phenotyping of mutants

Phototactic nectochaete larvae were transferred to a Petri dish and a single randomly selected larva was placed on a slide in 40 µl fNSW. The larva was scored for the startle response by touching it on the head with a tungsten needle. Only swimming larvae were assayed. For easier interpretation phenotyped larvae were classified into responders or non-responders. Single larvae were genotyped as described in the preceding section. The *PKD2-1^D137^* allele was detectable by gel electrophoresis, and thus only determined by sequencing in ambiguous cases.

### Calcium imaging

The *Palmi-3xHA-tdTomato-P2A-GCaMP6s* construct was assembled by restriction cloning from a plasmid kindly provided by Douglas Kim (*GCaMP6s*, Addgene plasmid #40753) and Martin Baier (*tdTomato*). *mRNA* was synthesized with the mMESSAGE mMACHINE™ T7 Transcription Kit (AM1344, Ambion). Animals injected with 2 µg/µl mRNA dissolved in RNAse-free water were tethered as described for the startle response quantification experiments, but using in this case a Ø5 cm glass-bottom dish in 5 ml volume. Tethered larvae were stimulated from the anterior with the filament set at a fixed value that invariably triggered the response. However,the maximum filament speed could not be directly measured for these experiments. Stimulus start was determined from the bright field channel (Video 4). In order to minimize muscle contractions, 500 µM Mecamylamine (M9020, Sigma) was added to the dish with the tethered larva. For calcium imaging in muscles no anesthetic compound was used. Slight X-Y shifts were corrected using descriptor-based series registration (Preibisch et al. 2010). Shifts in Z were accounted for with the tdTomato signal using the following formula:

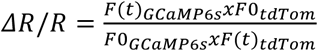

taken from (Böhm et al. 2016), where F0 is the average fluorescence prior to stimulation calculated from 1/2 the length of the pre-stimulation recorded period. Background signal (non-tissue signal) was used to adjust F0 and F(t). Most recordings were performed at 4 Hz, 3 recordings at 2.6 Hz.

### Circuit reconstruction

CR neurons and the downstream circuit were reconstructed from a ssTEM stack previously reported (Randel et al. 2015) using the collaborative annotation tool CATMAID (Saalfeld et al. 2009). Sensory endings in the left pygidial cirrus could not be imaged and thus CR neurons were not identified there. Synapses were defined as a discrete accumulation of vesicles in the inner side of the presynaptic membrane. The volume of each synapse was not considered (i.e. each synapse had a weight of 1, independently of how many sections it spanned), and thus the resulting network is a conservative representation of the synaptic strength between neurons. Any given neuron downstream of the CR neurons was included in the analysis if it had 3 or more synaptic contacts from at least 2 CR neurons. After applying this filter, additional neurons with only 1 or 2 synapses or only one upstream CR neuron were included if their bilateral counterparts (as defined by neuron morphology) were already part of the first selected set. The targets of the 1st layer in the network were likewise reconstructed, but only those belonging to the motorneuron class (i.e. cells innervating muscles or ciliary band cells) were included in the circuit. All neurons in the circuit were manually reviewed and weakly connected cells were double checked for accuracy of the synapse annotation. Eight fragments with three or more synapses downstream of the CR neurons but without a cell body were not included in the final circuit. A pair of glial cells, 4 sensory cells and two interneurons without a bilateral pair were also not included (Table 2 and 3). The additional penetrating ciliated sensory cells shown in Suppl. Figure 8 were reconstructed and reviewed only from sensory ending to cell body.

### Phylogenetic reconstruction

In addition to PKD1-1 and PKD2-1, a number of homologs to the PKD1 and TRPP families were found in the *Platynereis* transcriptome. A phylogenetic analysis was carried out in order to resolve their homology relationships. The amino acid sequences of the three human genes in the TRPP family (TRPP2,TRPP3, and TRPP5) and the five homologs of the PKD1 family (PKD1L1-3,PKDREJ and PC1) were used as queries in a BLAST search for homologs against the NCBI nr database. *Platynereis* PKD1 and PKD2 were also used as queries to find additional sequences. Care was taken to collect sequences from animals across the animal phylogeny. Additional sequences were obtained from the COMPAGEN (Hemmrich and Bosch 2008), and PlanMine (Brandl et al. 2015) databases. Full-length sequences were aligned using Clustal Omega (Sievers et al. 2014). For the joint PKD1-PKD2 tree, the alignment was cropped to span only the 6 transmembtane (TM) domains homologous to the TRP channel homology region common to both families. For the separate PKD2 and PKD1 family phylogenies any clearly alignable region was included. Only full sequences less than 90% identical were used for the alignment. GBlocks alignments (Talavera et al. 2007) were used for the phylogenetic reconstruction (Supp. Data 1). Maximum likelihood trees were recovered with PhyML (Guindon et al. 2010) using the SMS model selection tool (Lefort et al. 2017) and aLRT statistics (Anisimova et al. 2006).

### SEM of wild type and mutant nectochaete larvae

Mutant and age-matched wild type larvae were collected by phototaxis and simultaneously processed for SEM analysis. In brief, larvae were fixed in 3% Glutaraldehyde/PBS for 3 days (*PKD2-1^mut/mut^* and its wild type control) or for 1 month (*PKD1-1^i1/i1^* and its wild type control) at 4°C. Fixative was then washed out overnight in PBS, and samples were postfixed with 1% OsO4 for 2h on ice. Samples were gradually dehydrated with EtOH and critical point dried following standard protocols. Samples were sputter coated with an 8 nm layer of platinum (Bal-Tec MED010) and imaged with a Hitachi S-800 field emission scanning electron microscope at an accelerating voltage of 15KV. The genotype of fixed larvae was inferred from their sibling larvae.

### Predation experiments

Adult stages of male and female *C. typicus* were picked individually from culture bottles and transferred to a beaker with fNSW ca. 2h prior to the experiments. The experiments were carried out at 18°C in a Ø7 cm beaker completely wrapped with tin foil (i.e. dark conditions) and filled with 200 ml fNSW. 25 phototactic larvae of each genotype (25 age-matched wild type and 25 *PKD2-1^mut/mut^*) were transferred to the experimental container and then 5 or 10 copepods were added. Experiments were run for 12h or 24h without shaking (Almeda, van Someren Gréve, and Kiørboe 2017). At the end of the incubation period, live copepods were counted and every surviving larva was collected for genotyping. We used 12 mutant batches in 42 experiments (duplicates or triplicates with larvae from the same batch were simultaneously run for a given incubation time). In parallel to each experiment, a negative control with 25 larvae of each genotype but without copepods was run under the same environmental settings to assess mortality not related to predation. In the negative controls, 86% or more larvae survived, while the maximum survival rate with copepods was 70% (Table 1). Predation rates (*I*) for each genotype were calculated according to the following formula

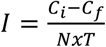

taken from (Almeda et al. 2017), where *C_i_* and *C_f_* are initial and final prey concentrations (prey/L), respectively, *N* is number of alive predators at the end of experiment and *T* is incubation time in days.

All videos were analyzed manually or with custom-written macros in FiJi, and data plots were generated in R. All panels were assembled into figures using Adobe Illustrator CS6 (Adobe Systems, Inc.).

## Acknowledgments

We thank Ada Kozlowska for help with the initial genotyping of *PKD1-1* mutants, Detlev Arendt for access to the *Platynereis* genome database, Nadine Randel for fixing samples for SEM, Dorothee Koch and Sinja Mattes for worm culture maintenance, Rocío Rodriguez for copepod culture maintenance and shipments, and Elizabeth Williams for comments on the manuscript. LABC was supported by a grant from the Deutsche Forschungsgemeinschaft (JE 777/3-1). RA was supported by a Marie Curie Intra-European fellowship (6240979) and by the Centre for Ocean Life, a VKR Center of Excellence funded by the VKR Foundation. The authors have no competing interests. All reagents and mutant stocks are available upon request.

## Supplementary Material

**Supplementary Figure 1:**
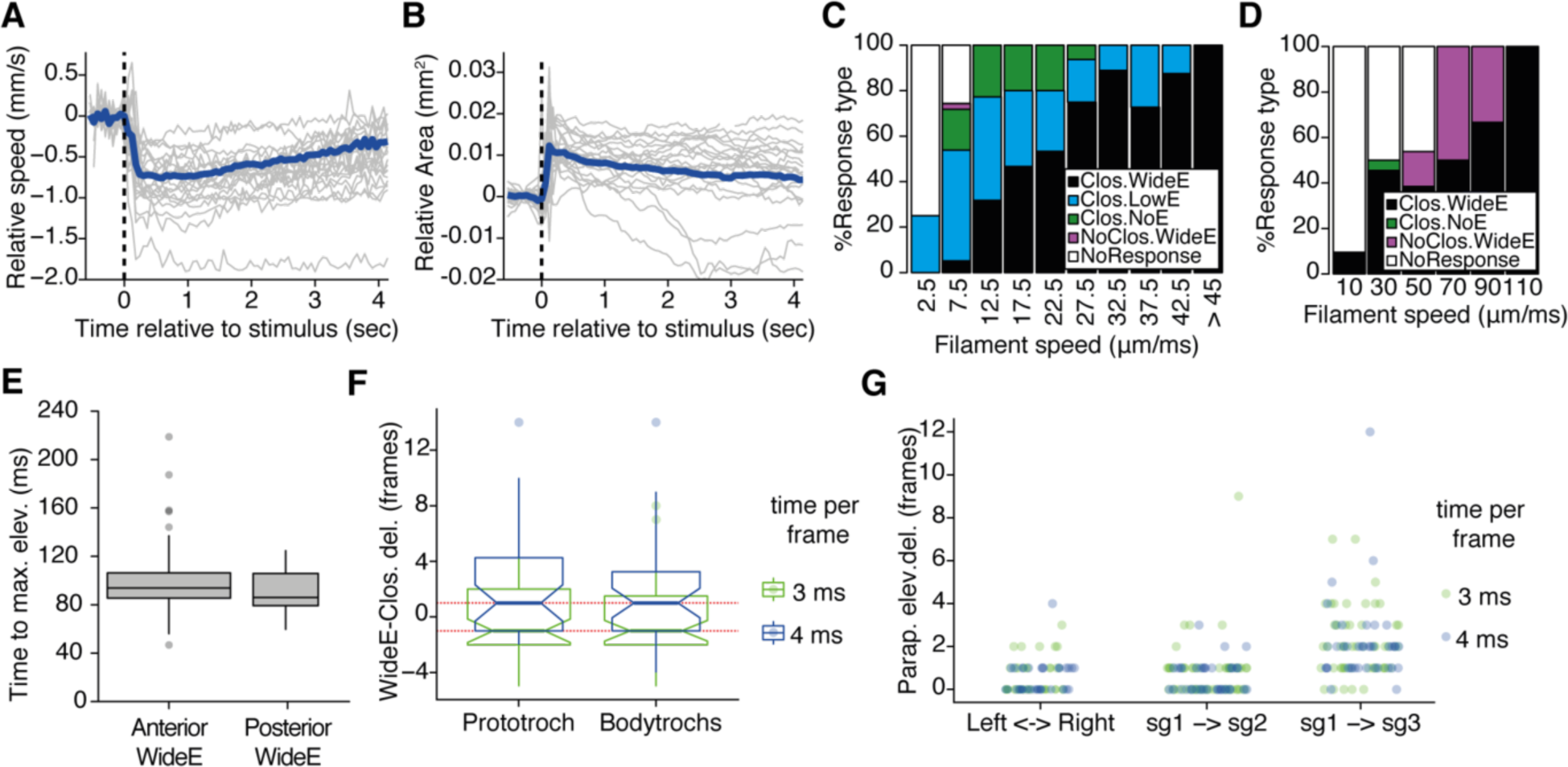
Descriptive statistics of the startle response in freely-swimming and tethered larvae. (**A-B**) Change in larval speed (B) or in area (B) upon vibration stimulus. Each value was normalized to average speed or average area prior to the stimulus. Blue lines represent the mean values and grey lines the individual values of 21 larvae measured from 3 batches. (**C-D**) Stacked barplot of startle response profiles upon head stimulation with a filament placed at 100 µm from the head (C) or pygidium (D). Midpoint values for each bin are shown in the abscissa. (**E**) Tukey boxplots of time from onset to maximum parapodial elevation for head and pygidial WideE responses. (**F**) Tukey boxplots of onset of WideE responses relative to prototroch or bodytroch ciliary arrests (i.e. WideE to closure delay) upon anterior stimulation. Red dashed lines mark the 1/−1 frame difference. (**G**) Frame difference between onset of left and right parapodial elevation in the 1^st^ segment (Left< −>Right), or the difference of the 1st relative to the 2nd parapodium, (sg1 − >sg2), or the 3rd parapodia (sg1−>sg3) on the same body side upon anterior stimulation. Notch in boxplots in F displays the 95% confidence interval around the median. Boxwidth in E and F is proportional to 
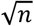
. Measurements in F and G are colored by recording speed. n=9 larvae in panels C-G, each tested multiple times.

**Supplementary Figure 2:**
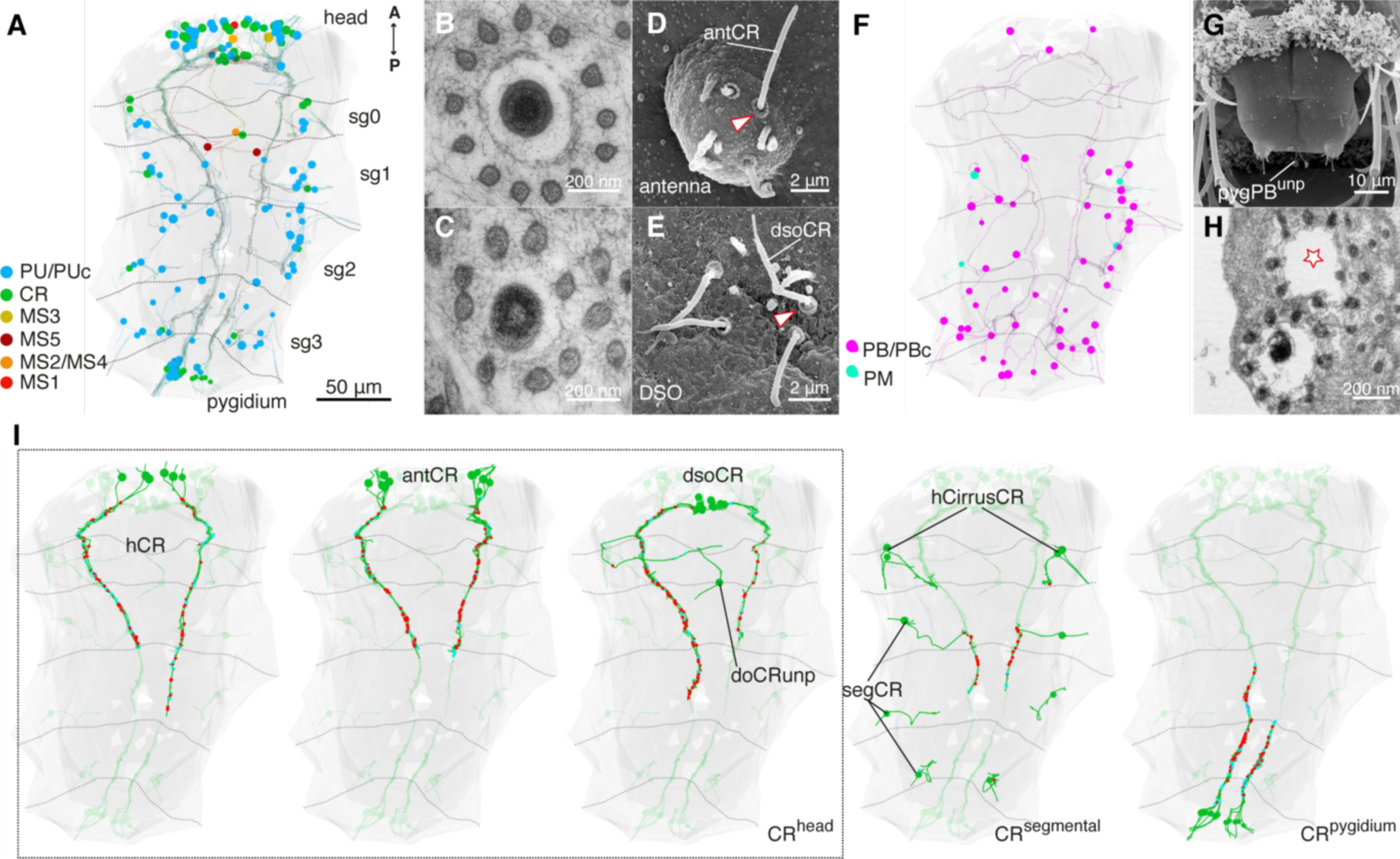
Penetrating ciliated sensory neurons in the nectochaete larva. (**A**) Penetrating uniciliated cell complement classified by sensory dendrite morphology. MS neurons are mostly asymmetric and localized to the head. CR and PU/PUc neurons are located around the body. (**B**) MS1 has a single cilium and collar of 13 microvilli. This sensory morphology is characteristic of MS neurons. (**C**) hPUc1 cilium with collar of 8 microvilli. PUc cells have irregularly shaped collars with variable numbers of thin microvilli. PU cells do not have a recognizable collar. (**D-E**) Organs with CR neurons. Antenna in D, and dorsal sensory organ (DSO) in E with multiple CR cells (antCR or dsoCR) recognized by the microvilli collar at the base (red arrowheads). (**F**)Penetrating biciliated (PB) and multiciliated (PM) cell complement. (**G**) Pygidium with multiple penetrating ciliated cells, including CR neurons and the single pygPB^unp^. (**H**) pygPB^unp^ has two cilia (one lost in this preparation, star) each with a collar of microvilli. (**I**) The 45 CR neurons (green) are localized to head, segments and pygidium. Output and input synapses are shown as red and cyan dots, respectively.

**Supplementary Figure 3:**
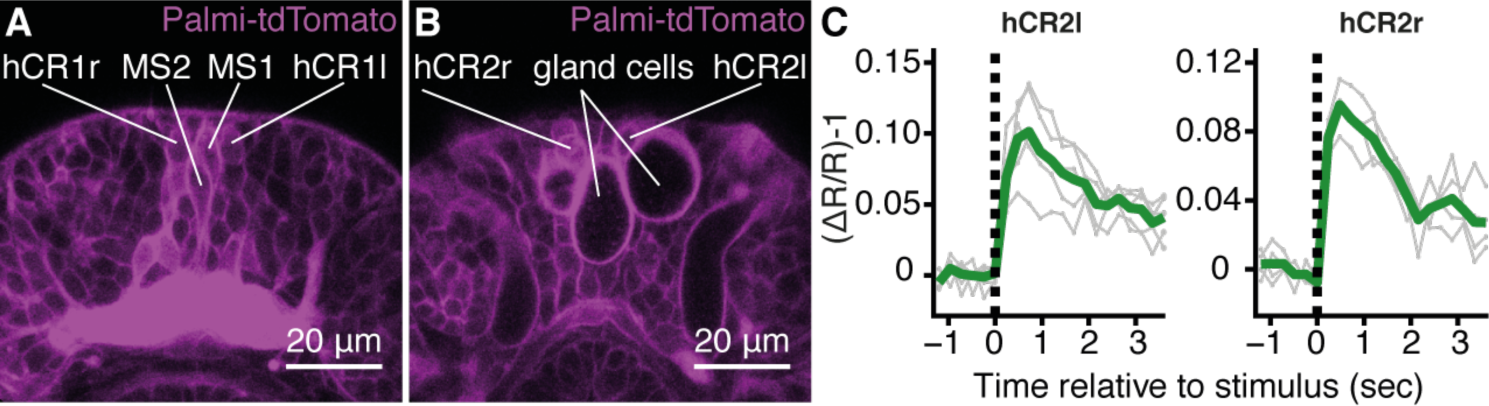
Distinct head CR and MS neurons are identifiable in living animals. (**A-B**) hCR1, hCR2 and MS1/2 cell bodies are identifiable in larvae injected with GCaMP6s-P2A-Palmi-tdTomato mRNA. hCR1 and MS1/2 are located in the middle of the head (A), while hCR2 cells are in a more ventral position and adjacent to the head gland cells (see Figure 2E). Only the signal from Palmi-tdtomato is shown. (**C-D**) Mean ratiometric fluorescence changes (green traces) of hCR2l, and hCR2r neurons upon stimulation with a vibrating filament placed 50m anterior to the head. 5 (hCR2l), and 4 (hCR2r) measurements (grey traces) from 2 animals are shown. The traces were aligned relative to the stimulus start (t=0).

**Supplementary Figure 4:**
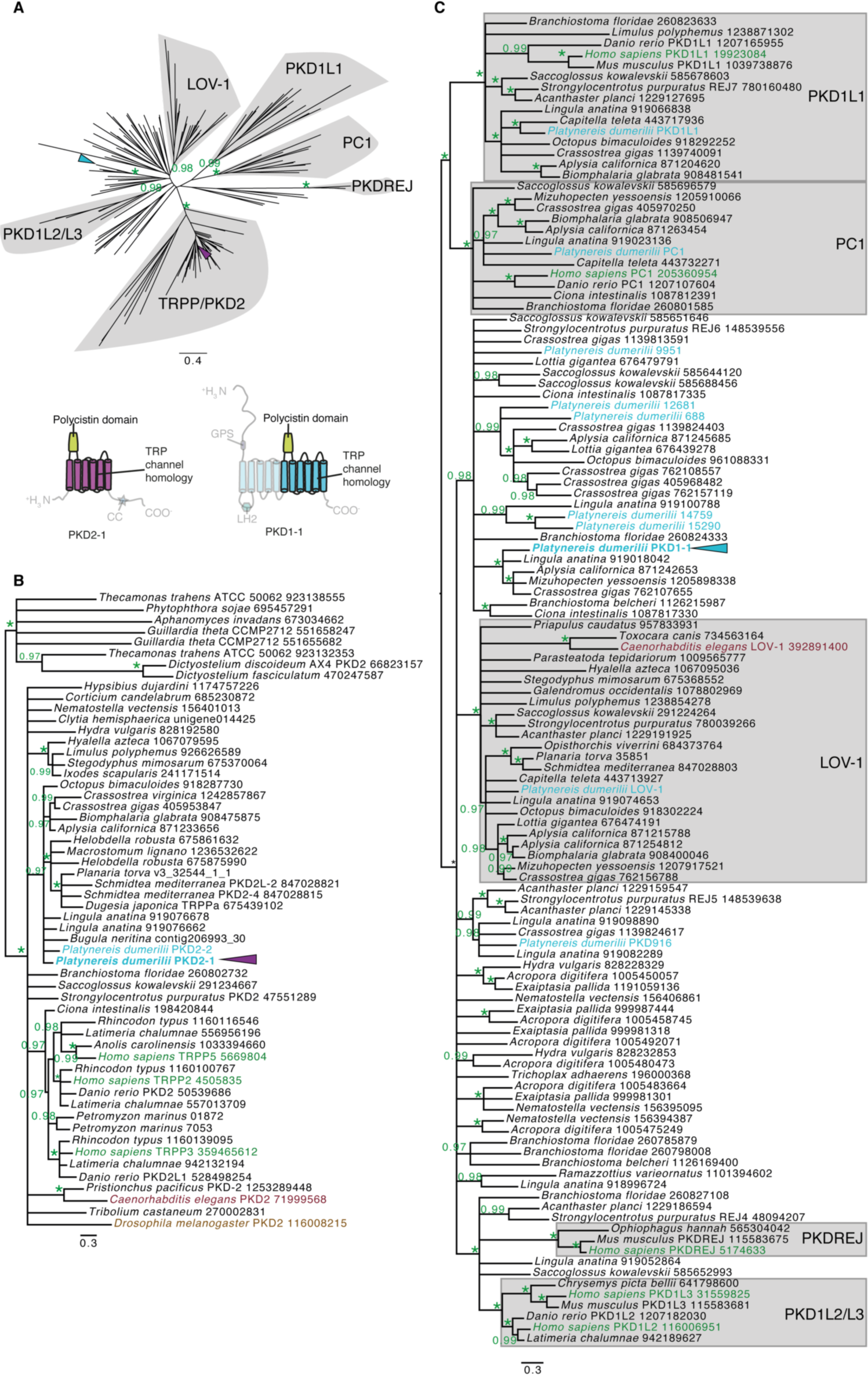
*Platynereis* PKD1-1 and PKD2-1 belong to the TRPP/PKD1 superfamily. (**A-C**) Maximum likelihood (ML) phylogenies of TRPP2 and/or PKD1 families. Well-supported families are outlined in gray color and named after experimentally characterized members. The location of *Platynereis* PKD1-1 and PKD2-1 in each tree is indicated with a cyan or a magenta arrowhead, respectively. Asterisks symbolize an aLRT value of 1. (**A**) ML phylogeny of the combined TRPP2 and PKD1 families (top) based on the common alignable region that includes the TRP channel homology domain and the poylcystin domain (highlighted in the PKD2-1 and PKD1-1 cartoon architectures at the bottom). (**B**-**C**) ML phylogeny of the TRPP/PKD2 family (B) or of the PKD1 family (C). Branching points with support values < 0.96 are displayed as a polytomy.

**Supplementary Figure 5:**
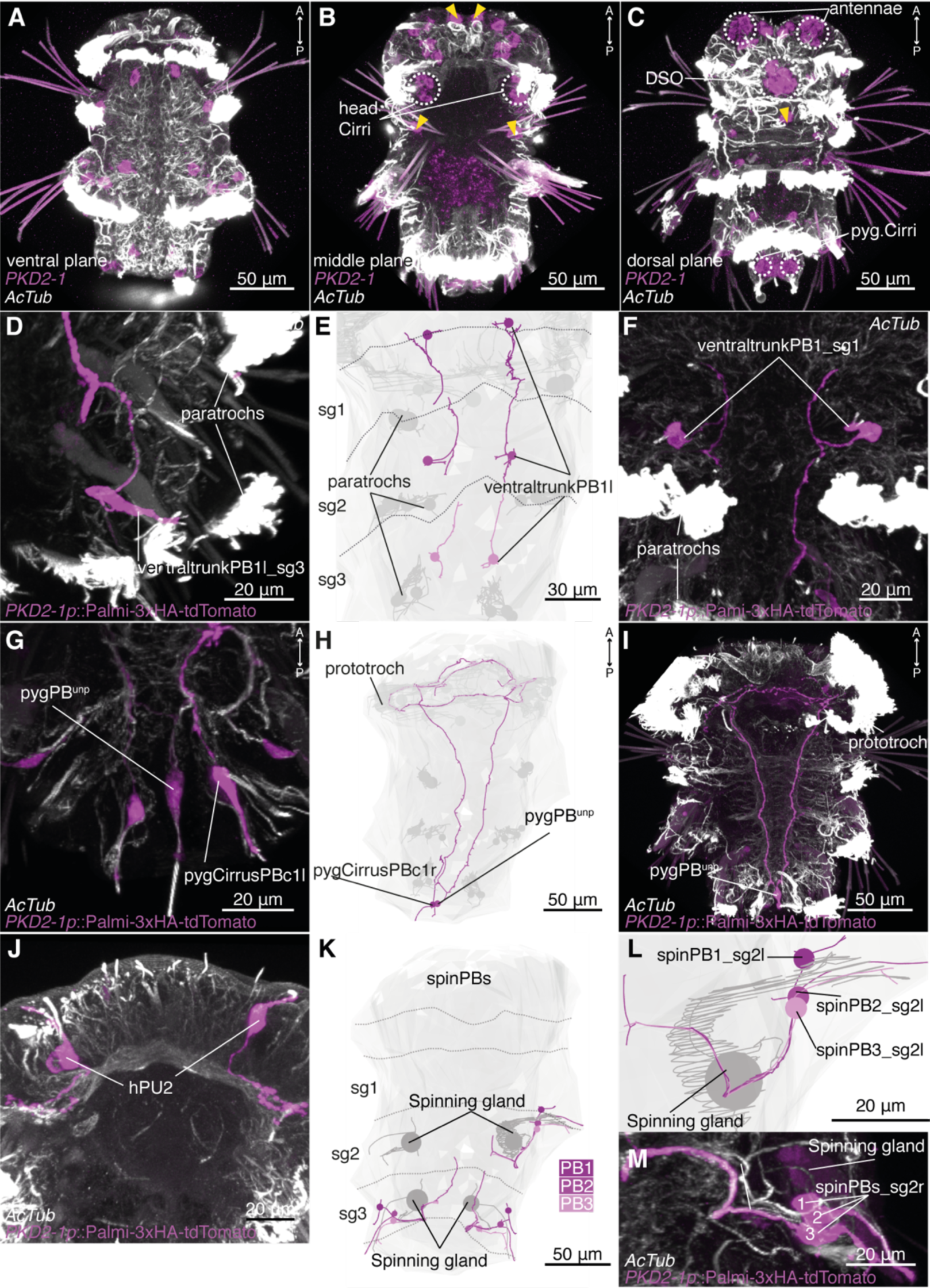
*PKD2-1* is expressed in CR and other penetrating ciliated neurons. (**A-C**) Different planes of a representative specimen showing expression of *PKD2-1* by whole mount *in situ* hybridisation. Yellow arrowheads point to putative CR neurons. White dotted circles demarcate sensory organs where CR neurons are present. Dorsal view in C. (**D**-**F**) *PKD2-1* is expressed in segmental biciliated cells in the trunk (ventraltrunkPBs) (D, F). The same cells were reconstructed in the EM volume (E). (**G**-**I**). *PKD2-1* is expressed in pygPB^unp^ (G, I), and in the single biciliated cell in the pygidial cirrus, pygCirrusPBc1 (G). H shows their corresponding EM reconstruction. (**J**) *PKD2-1* is expressed in hPU1 cells. EM reconstruction shown in Figure 2B. (**K-L**) *PKD2-1* is expressed in a group of three biciliated cells associated with the segmental spinning glands, spinPBs (M). Reconstruction of spinPB1-3 in each segment is shown in K (cells on the right body side in the 2nd segment could not be reconstructed). A close-up view is shown in L. Immunostainings against HA-Palmi-tdTomato expressed under the *PKD2-1* promoter are shown in D, F, G, I, J and M. Segment (sg) boundaries (dotted lines) in E and K were defined by muscle anatomy. Ventral views in all panels unless otherwise indicated.

**Supplementary Figure 6:**
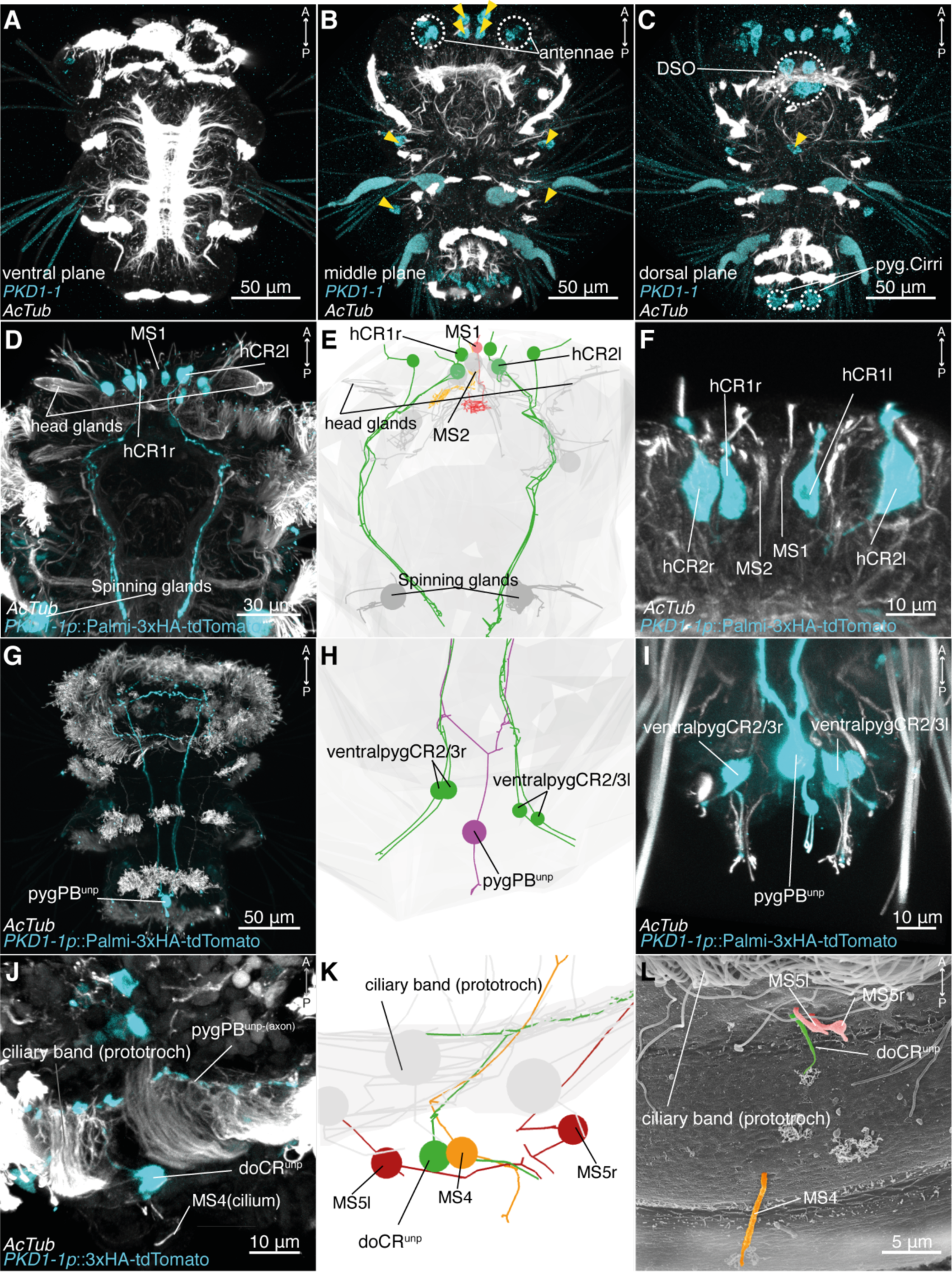
PKD1-1 is expressed in CR and pygPB^unp^ neurons. (**A-C**) Different planes of a representative specimen showing expression of PKD1-1 by whole mount in situ hybridization. Yellow arrowheads point to putative CR neurons. White dotted circles demarcate sensory organs where CR neurons are present. Dorsal view in C. (**D-F**) PKD1-1 is expressed in head CR neurons hCR1 and hCR2, but not in MS neurons MS1 and MS2 (D, F). The same cells were reconstructed in the EM volume (E). (**G-I**) PKD1-1 is expressed in pygPB^unp^ (G, I), and in CR neurons in the ventral side of the pygidium (ventralpygCR2/3) (I). H shows their corresponding EM reconstruction. (**J**) PKD1-1 is expressed in the unpaired dorsal CR neuron doCR^unp^ located slightly posterior to the prototroch. (**K**) Close-up view of doCR^unp^ reconstructed in the EM volume. Neighboring cells of the MS type (MS4 and MS5 cells) are also shown. (**L**) SEM view of the same region shown in K. Dorsal views in J-L. Cilia are colored according to cell colors in K. Immunostainings against HA-Palmi-tdTomato expressed under the PKD1-1 promoter are shown in D, F, G, I and J. Ventral views in all panels unless otherwise indicated.

**Supplementary Figure 7:**
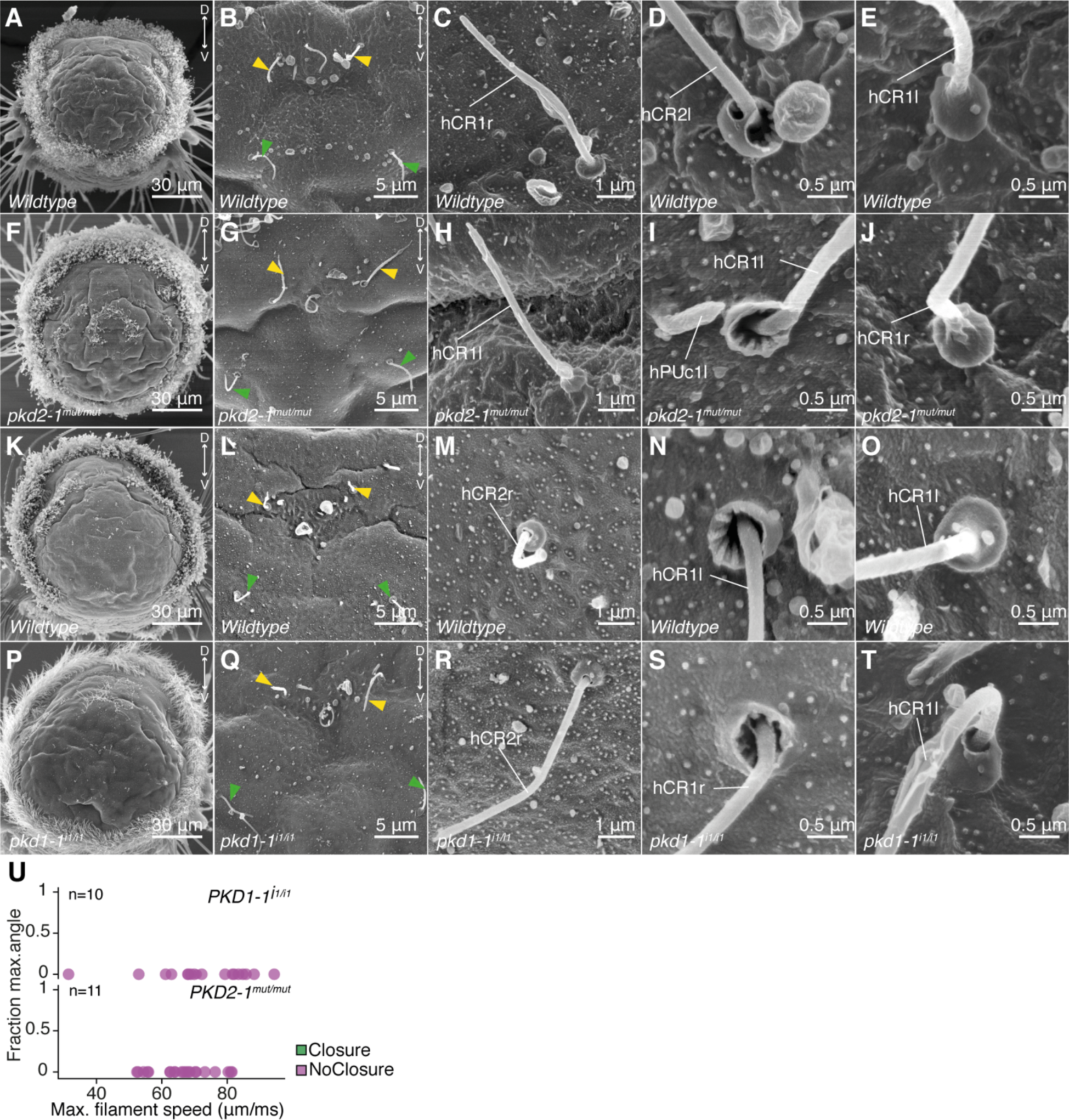
Cilia of hCR1 and hCR2 neurons in PKD1-1^i1/i1^ or PKD2-1^mut/mut^ larvae are morphologically similar to wildtype controls. (**A-J**). Morphology of hCR1 and hCR2 neurons in wildtype (A-E) and age-matched PKD2-1^mut/mut^ (F-J) larvae. (**K**-**T**). Morphology of hCR1 and hCR2 neurons in wildtype (K-O) and age-matched PKD1-1^i1/i1^ (P-T) larvae. hCR1 and hCR2 cilia are marked with yellow and green arrowheads, respectively, in B, G, L and Q. Examples of the “open” CR collar configuration are shown in D,I,N,S. Examples of the “closed” configuration are shown in E, J, O and T. (**U**) Parapodial elevation angles of PKD1-1 ^i1/i1^ or PKD2-1^mut/mut^ larvae as a function of filament speed. Stimulation filament ca. 100 µm from the pygidium.

**Supplementary Figure 8:**
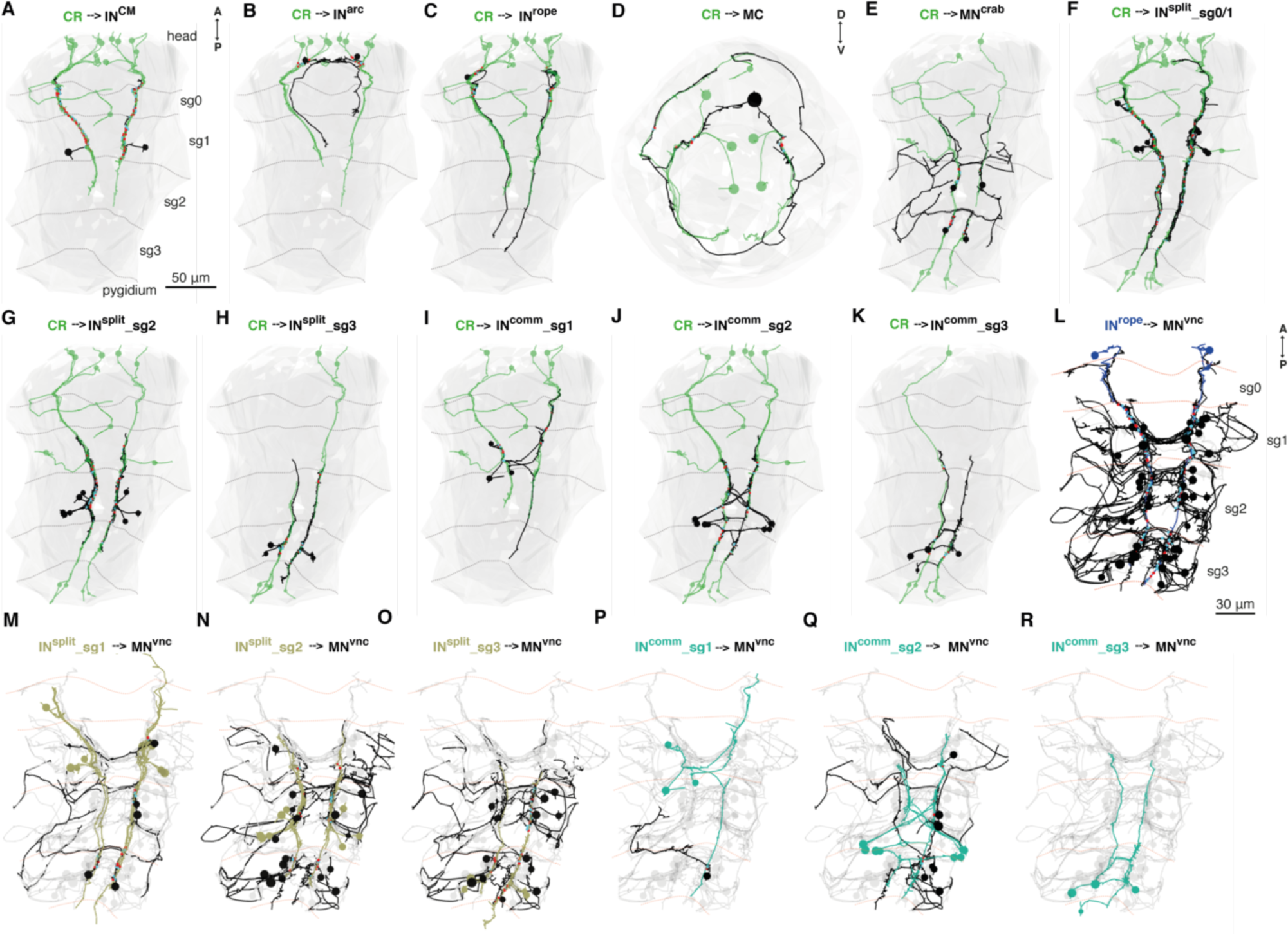
CR neurons target ipsilateral and commisural interneurons innervating VNC motorneurons. (**A-K**) Subsets of CR neurons (green cells) target diverse classes of interneurons and motorneurons (black). Only head CRs target CM, Arc, MC and Rope neurons (A-D), while both head and tail CR neurons target the segmentally arranged Split, Commissural and Crab neurons (E-K). Commissural interneurons have heterogeneous axonal projections. Apical view in D. (**L-R**) Interneurons downstream of CR neurons (colored) innervate distinct groups of VNC motorneurons (black). The complete set of (MN^vnc^) is colored in light gray. Ventral views in all panels except where indicated.

**Supplementary Figure 9:**
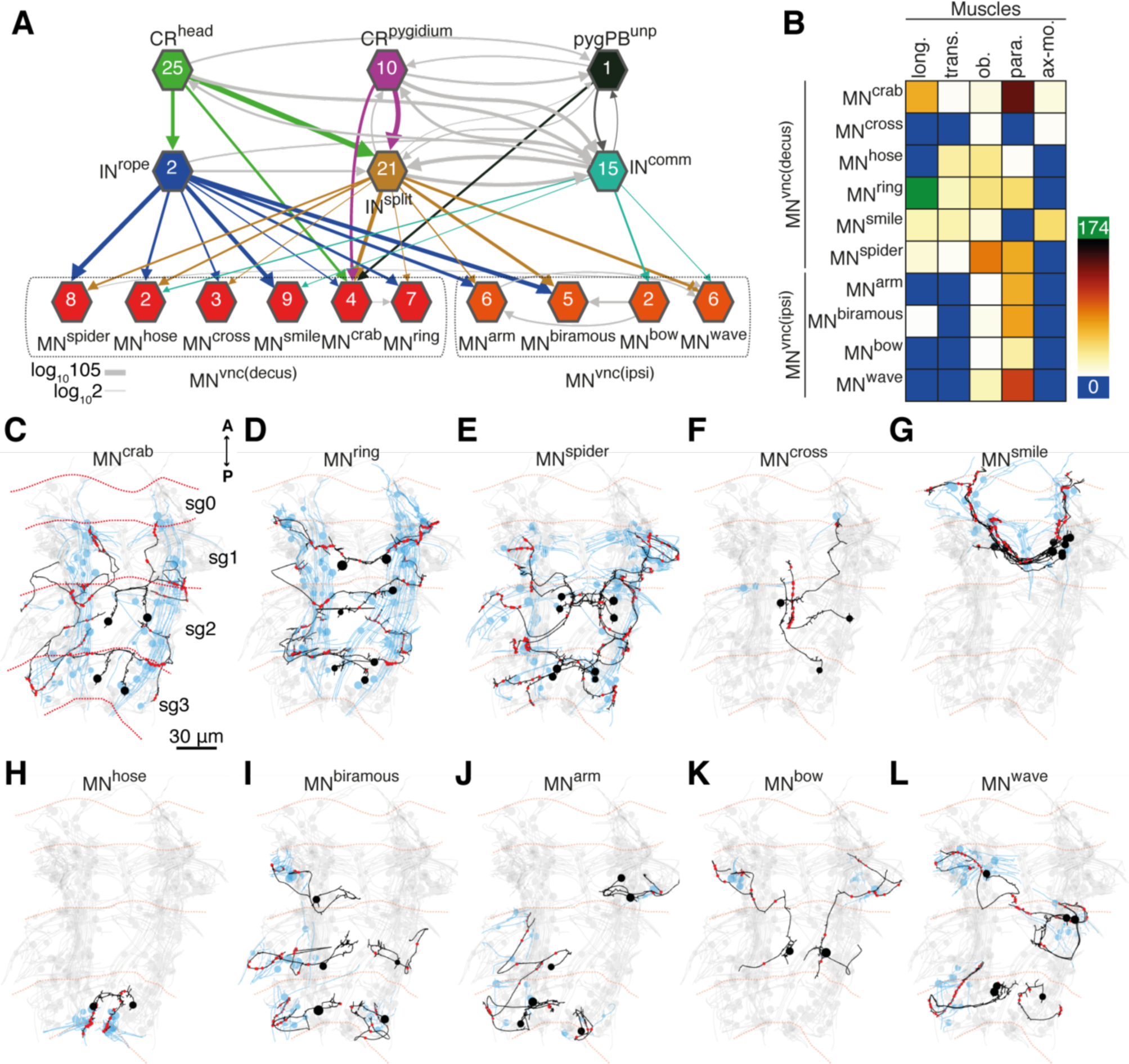
VNC motorneuron classes in the CR neural network. (**A**) Network from CR neurons to VNC motorneurons (MN^vnc^). Motorneurons are grouped by the 10 morphologically distinct types recognized and arranged by axonal projection patterns (decussating (MN^vnc(decuss)^ or ipsilateral MN^vnc(ipsi)^ (**B**) Connectivity matrix of Motorneuron to muscle class. (**C-L**) Reconstructed morphologies of the distinct motorneuron types (black) and the muscles they innervate (blue). As a reference, all the 273 muscle cells targeted by the whole set of motorneurons are shown and colored in light gray. Segment boundaries (sg) are indicated in C to L by dotted lines.

## References

1. Rodrigo Almeda, Hans van Someren Gréve, Thomas Kiørboe. Behavior is a major determinant of predation risk in zooplankton. Ecosphere 8, e01668 Wiley-Blackwell, 2017. <u>Link</u>

2. Maria Anisimova, Olivier Gascuel, Jack Sullivan. Approximate Likelihood-Ratio Test for Branches: A Fast Accurate, and Powerful lternative. Systematic Biology 55, 539–552 Oxford University Press (OUP), 2006. <u>Link</u>

3. Stephanie Bannister, Olga Antonova, Alessandra Polo, Claudia Lohs, Natalia Hallay, Agne Valinciute, Florian Raible, Kristin Tessmar-Raible. TALENs Mediate Efficient and Heritable Mutation of Endogenous Genes in the Marine Annelid Platynereis dumerilii. Genetics 197, 77–89 Genetics Society of America, 2014. <u>Link</u>

4. Maureen M. Barr, Paul W. Sternberg. A polycystic kidney-disease gene homologue required for male mating behaviour in C. elegans. Nature 401, 386–389 Springer Nature, 1999. <u>Link</u>

5. Maureen M. Barr, John DeModena, Douglas Braun, Can Q. Nguyen, David H. Hall, Paul W. Sternberg. The Caenorhabditis elegans autosomal dominant polycystic kidney disease gene homologs lov-1 and pkd-2 act in the same pathway. Current Biology 11, 1341–1346 Elsevier BV, 2001. <u>Link</u>

6. MM Barr, J DeModena, D Braun, CQ Nguyen, DH Hall, PW Sternberg. The Caenorhabditis elegans autosomal dominant polycystic kidney disease gene homologs lov-1 and pkd-2 act in the same pathway. Curr Biol 11, 1341–6 (2001).

7. MM Barr, PW Sternberg. A polycystic kidney-disease gene homologue required for male mating behaviour in C. elegans. Nature 401, 386–9 (1999).

8. Holger Brandl, HongKee Moon, Miquel Vila-Farré, Shang-Yun Liu, Ian Henry, Jochen C. Rink. PlanMine a mineable resource of planarian biology and biodiversity. Nucleic Acids Research 44, D764–D773 Oxford University Press (OUP), 2015. <u>Link</u>

9. Bernd-Ulrich Budelmann. Hydrodynamic Receptor Systems in Invertebrates. 607–631 In The Mechanosensory Lateral Line. Springer Nature, 1989. <u>Link</u>

10. Theodore Holmes Bullock. Comparative Neuroethology of Startle Rapid Escape, and Giant Fiber-Mediated Responses. 1–13 In Neural Mechanisms of Startle Behavior. Springer US, 1984. <u>Link</u>

11. Urs Lucas Böhm, Andrew Prendergast, Lydia Djenoune, Sophie Nunes Figueiredo, Johanna Gomez, Caleb Stokes, Sonya Kaiser, Maximilliano Suster, Koichi Kawakami, Marine Charpentier, Jean-Paul Concordet, Jean-Paul Rio, Filippo Del Bene, Claire Wyart. CSF-contacting neurons regulate locomotion by relaying mechanical stimuli to spinal circuits. Nature Communications 7, 10866 Springer Nature, 2016. <u>Link</u>

12. Albert Calbet, François Carlotti, Raymond Gaudy. The feeding ecology of the copepod Centropages typicus (Kröyer). Progress in Oceanography 72, 137–150 Elsevier BV, 2007. <u>Link</u>

13. Tsai-Wen Chen, Trevor J. Wardill, Yi Sun, Stefan R. Pulver, Sabine L. Renninger, Amy Baohan, Eric R. Schreiter, Rex A. Kerr, Michael B. Orger, Vivek Jayaraman, Loren L. Looger, Karel Svoboda, Douglas S. Kim. Ultrasensitive fluorescent proteins for imaging neuronal activity. Nature 499, 295–300 Springer Nature, 2013. <u>Link</u>

14. M Conzelmann, SL Offenburger, A Asadulina, T Keller, TA Münch, G Jékely. Neuropeptides regulate swimming depth of Platynereis larvae. Proc Natl Acad Sci U S A 108, E1174–83 (2011).

15. Bertrand Coste, Bailong Xiao, Jose S. Santos, Ruhma Syeda, Jörg Grandl, Kathryn S. Spencer, Sung Eun Kim, Manuela Schmidt, Jayanti Mathur, Adrienne E. Dubin, Mauricio Montal, Ardem Patapoutian. Piezo proteins are pore-forming subunits of mechanically activated channels. Nature 483, 176–181 Springer Nature, 2012. <u>Link</u>

16. Patrick Delmas, Bertrand Coste. Mechano-Gated Ion Channels in Sensory Systems. Cell 155, 278–284 Elsevier BV, 2013. <u>Link</u>

17. Alexandru S. Denes, Gáspár Jékely, Patrick R.H. Steinmetz, Florian Raible, Heidi Snyman, Benjamin Prudhomme, David E.K. Ferrier, Guillaume Balavoine, Detlev Arendt. Molecular Architecture of Annelid Nerve Cord Supports Common Origin of Nervous System Centralization in Bilateria. Cell 129, 277–288 Elsevier BV, 2007. <u>Link</u>

18. RC Eaton, RA Bombardieri, DL Meyer. The Mauthner-initiated startle response in teleost fish. J Exp Biol 66, 65–81 (1977).

19. Donald H. Edwards, William J. Heitler, Franklin B. Krasne. Fifty years of a command neuron: the neurobiology of escape behavior in the crayfish. Trends in Neurosciences 22, 153–161 Elsevier BV, 1999. <u>Link</u>

20. Antje HL Fischer, Thorsten Henrich, Detlev Arendt. The normal development of Platynereis dumerilii (Nereididae Annelida). Frontiers in Zoology 7, 31 Springer Nature, 2010. <u>Link</u>

21. A Fischer, A Dorresteijn. The polychaete Platynereis dumerilii (Annelida): a laboratory animal with spiralian cleavage, lifelong segment proliferation and a mixed benthic/pelagic life cycle. Bioessays 26, 314–25 (2004).

22. WO Friesen. Physiology of water motion detection in the medicinal leech. J Exp Biol 92, 255–75 (1981).

23. Stéphane Guindon, Jean-François Dufayard, Vincent Lefort, Maria Anisimova, Wim Hordijk, Olivier Gascuel. New Algorithms and Methods to Estimate Maximum-Likelihood Phylogenies: Assessing the Performance of PhyML 3.0. Systematic Biology 59, 307–321 Oxford University Press (OUP), 2010. <u>Link</u>

24. Martin Gühmann, Huiyong Jia, Nadine Randel, Csaba Verasztó, Luis A. Bezares-Calderón, Nico K. Michiels, Shozo Yokoyama, Gáspár Jékely. Spectral Tuning of Phototaxis by a Go-Opsin in the Rhabdomeric Eyes of Platynereis. Current Biology 25, 2265–2271 Elsevier BV, 2015. <u>Link</u>

25. ME Hale, HR Katz, MY Peek, RT Fremont. Neural circuits that drive startle behavior, with a focus on the Mauthner cells and spiral fiber neurons of fishes. J Neurogenet 30, 89–100 (2016).

26. K Hanaoka, F Qian, A Boletta, AK Bhunia, K Piontek, L Tsiokas, VP Sukhatme, WB Guggino, GG Germino. Co‐assembly of polycystin-1 and -2 produces unique cation-permeable currents. Nature 408, 990–4 (2000).

27. Georg Hemmrich, Thomas C.G. Bosch. Compagen a comparative genomics platform for early branching metazoan animals, reveals early origins of genes regulating stem-cell differentiation. BioEssays 30, 1010–1018 Wiley-Blackwell, 2008. <u>Link</u>

28. W. Charles Kerfoot. Combat between predatory copepods and their prey: Cyclops Epischura, and Bosmina. Limnology and Oceanography 23, 1089–1102 Wiley-Blackwell, 1978. <u>Link</u>

29. T Kiørboe, AW Visser. Predator and prey perception in copepods due to hydromechanical signals. Marine Ecology Progress Series 179, 81–95 Inter-Research Science Center, 1999. <u>Link</u>

30. Jin Hee Kim, Sang-Rok Lee, Li-Hua Li, Hye-Jeong Park, Jeong-Hoh Park, Kwang Youl Lee, Myeong-Kyu Kim, Boo Ahn Shin, Seok-Yong Choi. High Cleavage Efficiency of a 2A Peptide Derived from Porcine Teschovirus-1 in Human Cell Lines Zebrafish and Mice. PLoS ONE 6, e18556 Public Library of Science (PLoS), 2011. <u>Link</u>

31. M.F. Knapp, P.J. Mill. The fine structure of ciliated sensory cells in the epidermis of the earthworm Lumbricus terrestris. Tissue and Cell 3, 623–636 Elsevier BV, 1971. <u>Link</u>

32. H Korn, DS Faber. The Mauthner cell half a century later: a neurobiological model for decision-making? Neuron 47, 13–28 (2005).

33. Alix M.B. Lacoste, David Schoppik, Drew N. Robson, Martin Haesemeyer, Ruben Portugues, Jennifer M. Li, Owen Randlett, Caroline L. Wee, Florian Engert, Alexander F. Schier. A Convergent and Essential Interneuron Pathway for Mauthner-Cell-Mediated Escapes. Current Biology 25, 1526–1534 Elsevier BV, 2015. <u>Link</u>

34. A. Lauri, T. Brunet, M. Handberg-Thorsager, A. H. L. Fischer, O. Simakov, P. R. H. Steinmetz, R. Tomer, P. J. Keller, D. Arendt. Development of the annelid axochord: Insights into notochord evolution. Science 345, 1365–1368 American Association for the Advancement of Science (AAAS), 2014. <u>Link</u>

35. Vincent Lefort, Jean-Emmanuel Longueville, Olivier Gascuel. SMS: Smart Model Selection in PhyML. Molecular Biology and Evolution 34, 2422–2424 Oxford University Press (OUP), 2017. <u>Link</u>

36. S Lin, BT Staahl, RK Alla, JA Doudna. Enhanced homology-directed human genome engineering by controlled timing of CRISPR/Cas9 delivery. Elife 3, e04766 (2014).

37. G Mackie, R Meech. Central circuitry in the jellyfish Aglantha. I: The relay system. J Exp Biol 198, 2261–70 (1995a).

38. G Mackie, R Meech. Central circuitry in the jellyfish Aglantha. II: The ring giant and carrier systems. J Exp Biol 198, 2271–8 (1995b).

39. GO Mackie, CL Singla, C Thiriot-Quievreux. Nervous control of ciliary activity in gastropod larvae. Biol Bull 151, 182–99 (1976).

40. George O. Mackie, C. L. Singla, Catherine Thiriot-Quievreux. Nervous control of ciliary activity in gastropod larvae. The Biological Bulletin 151, 182–199 University of Chicago Press, 1976. <u>Link</u>

41. SM Nauli, FJ Alenghat, Y Luo, E Williams, P Vassilev, X Li, AE Elia, W Lu, EM Brown, SJ Quinn, DE Ingber, J Zhou. Polycystins 1 and 2 mediate mechanosensation in the primary cilium of kidney cells. Nat Genet 33, 129–37 (2003).

42. Surya M. Nauli, Francis J. Alenghat, Ying Luo, Eric Williams, Peter Vassilev, Xiaogang Li, Andrew E. H. Elia, Weining Lu, Edward M. Brown, Stephen J. Quinn, Donald E. Ingber, Jing Zhou. Polycystins 1 and 2 mediate mechanosensation in the primary cilium of kidney cells. Nature Genetics 33, 129–137 Springer Nature, 2003. <u>Link</u>

43. T Ohyama, CM Schneider-Mizell, RD Fetter, JV Aleman, R Franconville, M Rivera-Alba, BD Mensh, KM Branson, JH Simpson, JW Truman, A Cardona, M Zlatic. A multilevel multimodal circuit enhances action selection in Drosophila. Nature 520, 633–9 (2015).

44. J. Timothy Pennington, Fu-Shiang Chia. Morphological and behavioral defenses of trochophore larvae of Sabellaria cementarium (Polychaeta) against four planktonic predators. The Biological Bulletin 167, 168–175 University of Chicago Press, 1984. <u>Link</u>

45. Christine E. Phillips, W. Otto Friesen. Ultrastructure of the water-movement-sensitive sensilla in the medicinal leech. Journal of Neurobiology 13, 473–486 Wiley-Blackwell, 1982. <u>Link</u>

46. Stephan Preibisch, Stephan Saalfeld, Johannes Schindelin, Pavel Tomancak. Software for bead-based registration of selective plane illumination microscopy data. Nature Methods 7, 418–419 Springer Nature, 2010. <u>Link</u>

47. Günter Purschke. Sense organs in polychaetes (Annelida). Hydrobiologia 535-536, 53–78 Springer Nature, 2005. <u>Link</u>

48. Günter Purschke, Maja Hugenschütt, Lisa Ohlmeyer, Heiko Meyer, Dirk Weihrauch. Structural analysis of the branchiae and dorsal cirri in Eurythoe complanata (Annelida Amphinomida). Zoomorphology 136, 1–18 Springer Nature, 2016. <u>Link</u>

49. Nadine Randel, Réza Shahidi, Csaba Verasztó, Luis A Bezares-Calderón, Steffen Schmidt, Gáspár Jékely. Inter-individual stereotypy of thePlatynereislarval visual connectome. eLife 4 eLife Sciences Organisation Ltd., 2015. <u>Link</u>

50. Nadine Randel, Albina Asadulina, Luis A Bezares-Calderón, Csaba Verasztó, Elizabeth A Williams, Markus Conzelmann, Réza Shahidi, Gáspár Jékely. Neuronal connectome of a sensory-motor circuit for visual navigation. eLife 3 eLife Sciences Organisation Ltd., 2014. <u>Link</u>

51. Nadine Randel, Réza Shahidi, Csaba Verasztó, Luis A Bezares-Calderón, Steffen Schmidt, Gáspár Jékely. Inter-individual stereotypy of the Platynereis larval visual connectome. eLife 4 eLife Sciences Organisation Ltd., 2015. <u>Link</u>

52. N Randel, A Asadulina, LA Bezares-Calderón, C Verasztó, EA Williams, M Conzelmann, R Shahidi, G Jékely. Neuronal connectome of a sensory-motor circuit for visual navigation. Elife 3 (2014).

53. A Roberts, GO Mackie. The giant axon escape system of a hydrozoan medusa, Aglantha digitale. J Exp Biol 84, 303–18 (1980).

54. Kerrianne Ryan, Zhiyuan Lu, Ian A. Meinertzhagen. Circuit Homology between Decussating Pathways in the Ciona Larval CNS and the Vertebrate Startle-Response Pathway. Current Biology 27, 721–728 Elsevier BV, 2017. <u>Link</u>

55. S. Saalfeld, A. Cardona, V. Hartenstein, P. Tomancak. CATMAID: collaborative annotation toolkit for massive amounts of image data. Bioinformatics 25, 1984–1986 Oxford University Press (OUP), 2009. <u>Link</u>

56. J. D. Sander, P. Zaback, J. K. Joung, D. F. Voytas, D. Dobbs. Zinc Finger Targeter (ZiFiT): an engineered zinc finger/target site design tool. Nucleic Acids Research 35, W599–W605 Oxford University Press (OUP), 2007. <u>Link</u>

57. A. Schlawny, C. Grünig, H. -D. Pfannenstiel. Sensory and secretory cells ofOphryotrocha puerilis (Polychaeta). Zoomorphology 110, 209–215 Springer Nature, 1991. <u>Link</u>

58. Réza Shahidi, Elizabeth A Williams, Markus Conzelmann, Albina Asadulina, Csaba Verasztó, Sanja Jasek, Luis A Bezares-Calderón, Gáspár Jékely. A serial multiplex immunogold labeling method for identifying peptidergic neurons in connectomes. eLife 4 eLife Sciences Organisation Ltd., 2015. <u>Link</u>

59. Reza Sharif-Naeini, Joost H.A. Folgering, Delphine Bichet, Fabrice Duprat, Inger Lauritzen, Malika Arhatte, Martine Jodar, Alexandra Dedman, Franck C. Chatelain, Uwe Schulte, Kevin Retailleau, Laurent Loufrani, Amanda Patel, Frederick Sachs, Patrick Delmas, Dorien J.M. Peters, Eric Honoré. Polycystin-1 and -2 Dosage Regulates Pressure Sensing. Cell 139, 587–596 Elsevier BV, 2009. <u>Link</u>

60. Peter S. Shen, Xiaoyong Yang, Paul G. DeCaen, Xiaowen Liu, David Bulkley, David E. Clapham, Erhu Cao. The Structure of the Polycystic Kidney Disease Channel PKD2 in Lipid Nanodiscs. Cell 167, 763–773.e11 Elsevier BV, 2016. <u>Link</u>

61. F. Sievers, A. Wilm, D. Dineen, T. J. Gibson, K. Karplus, W. Li, R. Lopez, H. McWilliam, M. Remmert, J. Soding, J. D. Thompson, D. G. Higgins. Fast scalable generation of high-quality protein multiple sequence alignments using Clustal Omega. Molecular Systems Biology 7, 539–539 Wiley-Blackwell, 2014. <u>Link</u>

62. J.N Sigger, D.A Dorsett. Physiological and anatomical relationships of the 4B motoneurons of Nereis. Comparative Biochemistry and Physiology Part A: Physiology 84, 783–788 Elsevier BV, 1986. <u>Link</u>

63. J. E. Smith. The Nervous Anatomy of the Body Segments of Nereid Polychaetes. Philosophical Transactions of the Royal Society B: Biological Sciences 240, 135–196 The Royal Society, 1957. <u>Link</u>

64. K. A. Steigelman, A. Lelli, X. Wu, J. Gao, S. Lin, K. Piontek, C. Wodarczyk, A. Boletta, H. Kim, F. Qian, G. Germino, G. S. G. Geleoc, J. R. Holt, J. Zuo. Polycystin-1 Is Required for Stereocilia Structure But Not for Mechanotransduction in Inner Ear Hair Cells. Journal of Neuroscience 31, 12241–12250 Society for Neuroscience, 2011. <u>Link</u>

65. Suguru Takagi, Benjamin Thomas Cocanougher, Sawako Niki, Dohjin Miyamoto, Hiroshi Kohsaka, Hokto Kazama, Richard Doty Fetter, James William Truman, Marta Zlatic, Albert Cardona, Akinao Nose. Divergent Connectivity of Homologous Command-like Neurons Mediates Segment-Specific Touch Responses in Drosophila. Neuron 96, 1373–1387.e6 Elsevier BV, 2017. <u>Link</u>

66. Gerard Talavera, Jose Castresana, Karl Kjer, Rod Page, Jack Sullivan. Improvement of Phylogenies after Removing Divergent and Ambiguously Aligned Blocks from Protein Sequence Alignments. Systematic Biology 56, 564–577 Oxford University Press (OUP), 2007. <u>Link</u>

67. Daniel Thiel, Philipp Bauknecht, Gáspár Jékely, Andreas Hejnol. An ancient FMRFamide-related peptidereceptor pair induces defence behaviour in a brachiopod larva. Open Biology 7, 170136 The Royal Society, 2017. <u>Link</u>

68. Raju Tomer, Alexandru S. Denes, Kristin Tessmar-Raible, Detlev Arendt. Profiling by Image Registration Reveals Common Origin of Annelid Mushroom Bodies and Vertebrate Pallium. Cell 142, 800–809 Elsevier BV, 2010. <u>Link</u>

69. Csaba Verasztó, Nobuo Ueda, Luis A Bezares-Calderón, Aurora Panzera, Elizabeth A Williams, Réza Shahidi, Gáspár Jékely. Ciliomotor circuitry underlying whole-body coordination of ciliary activity in the Platynereis larva. eLife 6 eLife Sciences Organisation Ltd., 2017. <u>Link</u>

70. C Verasztó, N Ueda, LA Bezares-Calderón, A Panzera, EA Williams, R Shahidi, G Jékely. Ciliomotor circuitry underlying whole-body coordination of ciliary activity in the Platynereis larva. Elife 6 (2017).

71. EA Williams, C Verasztó, S Jasek, M Conzelmann, R Shahidi, P Bauknecht, O Mirabeau, G Jékely. Synaptic and peptidergic connectome of a neurosecretory center in the annelid brain. Elife 6 (2017).

72. Reinhard Windoffer, Wilfried Westheide. The Nervous System of the Male Dinophilus gyrociliatus (Annelida: Polychaeta). I. Number Types and Distribution Pattern of Sensory Cells. Acta Zoologica 69, 55–64 Wiley-Blackwell, 1988. <u>Link</u>

73. S. Yoshiba, H. Shiratori, I. Y. Kuo, A. Kawasumi, K. Shinohara, S. Nonaka, Y. Asai, G. Sasaki, J. A. Belo, H. Sasaki, J. Nakai, B. Dworniczak, B. E. Ehrlich, P. Pennekamp, H. Hamada. Cilia at the Node of Mouse Embryos Sense Fluid Flow for Left-Right Determination via Pkd2. Science 338, 226–231 American Association for the Advancement of Science (AAAS), 2012. <u>Link</u>

74. R S Zucker. Crayfish escape behavior and central synapses. I. Neural circuit exciting lateral giant fiber. Journal of Neurophysiology 35, 599–620 American Physiological Society, 1972. <u>Link</u>

